# Graded and sharp transitions in semantic function in left temporal lobe

**DOI:** 10.1101/2023.05.04.539459

**Authors:** Katya Krieger-Redwood, Xiuyi Wang, Nicholas Souter, Tirso Rene del Jesus Gonzalez Alam, Jonathan Smallwood, Rebecca L. Jackson, Elizabeth Jefferies

## Abstract

Recent work has focussed on how patterns of functional change within the temporal lobe relate to whole-brain dimensions of intrinsic connectivity variation (Margulies et al., 2016). We examined two such ‘connectivity gradients’ reflecting the separation of (i) unimodal versus heteromodal and (ii) visual versus auditory-motor cortex, examining visually presented verbal associative and feature judgments, plus picture-based context and emotion generation. Functional responses along the first dimension sometimes showed graded change between modality-tuned and heteromodal cortex (in the verbal matching task), and other times showed sharp functional transitions, with deactivation at the extremes and activation in the middle of this gradient (internal generation). The second gradient revealed more visual than auditory-motor activation, regardless of content (associative, feature, context, emotion) or task process (matching/generation). We also uncovered subtle differences across each gradient for content type, which predominantly manifested as differences in relative magnitude of activation or deactivation.

## 1. Introduction

The temporal lobes are associated with multimodal conceptual processing, but their functional organisation remains controversial. While most researchers now agree that temporal cortex supports aspects of semantic processing, there remains uncertainty about whether site(s) within this broad region support multimodal concepts across categories (Lambon Ralph, Jefferies, Patterson, & Rogers, 2017), or whether there are distinct regions that underpin specific aspects of knowledge pertaining to words, objects, scenes, emotions, social cognition (e.g., theory of mind, ToM) and people (Hoenig, Sim, Bochev, Herrnberger, & Kiefer, 2008; Martin, 2007; Persichetti, Denning, Gotts, & Martin, 2021; Simmons & Martin, 2009; Simmons, Martin, & Barsalou, 2005). There is ongoing debate about whether the organisation of temporal cortex reflects a patchwork of functions (Malone, Glezer, Kim, Jiang, & Riesenhuber, 2016; Olson, Plotzker, & Ezzyat, 2007; Persichetti et al., 2021) or whether there is graded functional change such that subregions of this brain area show systematic functional transitions based on their location (Binney, Hoffman, & Lambon Ralph, 2016; Lambon Ralph et al., 2017; Visser, Jefferies, Embleton, & Lambon Ralph, 2012; Visser & Lambon Ralph, 2011).

Much of what we know about the organisation of temporal cortex comes from neuropsychological studies of semantic dementia (characterised by progressive neurodegeneration of anterior and inferior temporal cortex accompanied by degradation of conceptual knowledge) and meta-analyses of neuroimaging studies (Binder, Desai, Graves, & Conant, 2009; Hung, Wang, Wang, & Bi, 2020; Jefferies & Lambon Ralph, 2006; Lambon Ralph, Cipolotti, Manes, & Patterson, 2010; Rice, Lambon Ralph, & Hoffman, 2015; Snowden et al., 2001; Snowden, Thompson, & Neary, 2004; Visser, Jefferies, & Lambon Ralph, 2009; Warrington, 1975; Warrington & Cipolotti, 1996). Both of these methods lack spatial resolution and are therefore not ideal for resolving questions such as which regions in temporal cortex are crucial for aspects of long-term conceptual representation (De Panfilis & Schwarzbauer, 2005; Devlin et al., 2000).

Functional magnetic resonance imaging (fMRI) investigations of anterior and medial portions of temporal cortex have been hampered by magnetic susceptibility artefacts caused by the proximity of this brain region to the air-filled sinuses, which produces signal loss and distortions. For this reason, positron emission tomography studies have often recovered anterior temporal lobe responses in semantic tasks while fMRI studies have not (Martin, Wiggs, Ungerleider, & Haxby, 1996; Visser et al., 2009). Questions remain about how these responses relate to the organisation of semantic memory (Davis & Yee, 2021; Lambon Ralph et al., 2017; Simmons & Martin, 2009). More recently fMRI sequences designed to recover better signal from anterior and medial temporal cortex have led to an increase in our understanding of this region. These sequences utilise a smaller echo time and/or combine this with multiple echoes; for example, modern multiband multi-echo (MBME) sequences provide better signal-to-noise in these regions (Embleton, Haroon, Morris, Lambon Ralph, & Parker, 2010; Halai, Parkes, & Welbourne, 2015; Halai, Welbourne, Embleton, & Parkes, 2014; Poser & Norris, 2007; Poser, Versluis, Hoogduin, & Norris, 2006). Several recent studies using these imaging sequences have examined the functional organisation of temporal cortex in semantic cognition (e.g., Balgova, Diveica, Walbrin, & Binney, 2022; Binney, Embleton, Jefferies, Parker, & Lambon Ralph, 2010; Binney et al., 2016; Ovando-Tellez et al., 2022; Persichetti et al., 2021; Rice, Hoffman, Binney, & Lambon Ralph, 2018; Visser, Embleton, Jefferies, Parker, & Ralph, 2010). Ventral anterior temporal cortex has emerged as a potential multi-modal conceptual integrator due to both its activation profile and structural connectivity – with long range connectivity from primary sensory areas gradually converging on the ventral anterior temporal cortex (Bajada et al., 2017; Chen et al., 2016; Jackson, Bajada, Rice, Cloutman, & Lambon Ralph, 2018; Shimotake et al., 2015); however, this interpretation is contested by other research groups who also used modern imaging techniques to conduct parcellations but with different analysis strategies, and argued for a patchwork organisation with no integrative convergence zone in the temporal lobe (Persichetti et al., 2021).

While the debate continues regarding the organisation of semantic memory, most theories agree that the representation of concepts requires a coordinated response across the cortex (Barsalou, 2008; Barsalou, 2016; Binder & Desai, 2011; Davis & Yee, 2021; Jefferies, 2013; Martin, 2016). It is also widely accepted that the temporal lobe has an important role in semantic cognition, but the function and organisation is still debated. For example, the temporal lobe could be organised into a patchwork of functional specificity (e.g., Persichetti et al., 2021), or as a gradient of function converging on an amodal ‘hub’ (the graded hub account; e.g., Lambon Ralph et al., 2017; Rogers et al., 2021). Both theoretical viewpoints find support in studies that reveal activation in temporal cortex for a wide range of semantic concepts, but debate the underlying functional organisation, and tend to consider activation at different spatial scales, from fine-grained parcels to broader patterns of semantic activation. Yet the importance of long-range connections across the brain for conceptual processing is consistent across theoretical perspectives (Bajada et al., 2017; Jackson et al., 2018; Lambon Ralph et al., 2017; Martin, 2016; Persichetti et al., 2021): both views agree that functional organisation should reflect connectivity.

Recent work has focussed on how patterns of functional change within the temporal lobe relate to whole-brain connectivity gradients, which capture systematic changes in intrinsic connectivity across the cortical surface (Margulies et al., 2016; Wang, Margulies, Smallwood, & Jefferies, 2020). Rather than focusing on discrete regions, these gradients are able to capture how the brain works in a coordinated fashion. The first gradient (G1) in whole-brain decompositions of resting-state fMRI captures the difference in connectivity between unimodal and heteromodal cortex (Figure 1), with the heteromodal end of this dimension associated with aspects of cognition that are guided by memory and might involve more abstract codes (Murphy et al., 2018; Murphy et al., 2017; Murphy et al., 2019; Smallwood et al., 2021). The second component captured by these decompositions of intrinsic connectivity (G2) reflects the separation between sensorimotor domains, with auditory regions on one end and visual systems on the other (Figure 1); therefore, while the first gradient might relate to the way in which semantic processing draws on sensorimotor and heteromodal processes (in a binary or graded fashion), G2 could capture the degree to which different tasks leverage these sensorimotor features. For example, portions of G2 closer to auditory systems might activate more for verbal semantic material (regardless of input domain), while regions at the other end of G2 (closer to vision) might preferentially activate for visually-instantiated concrete concepts.

**Figure 1.**
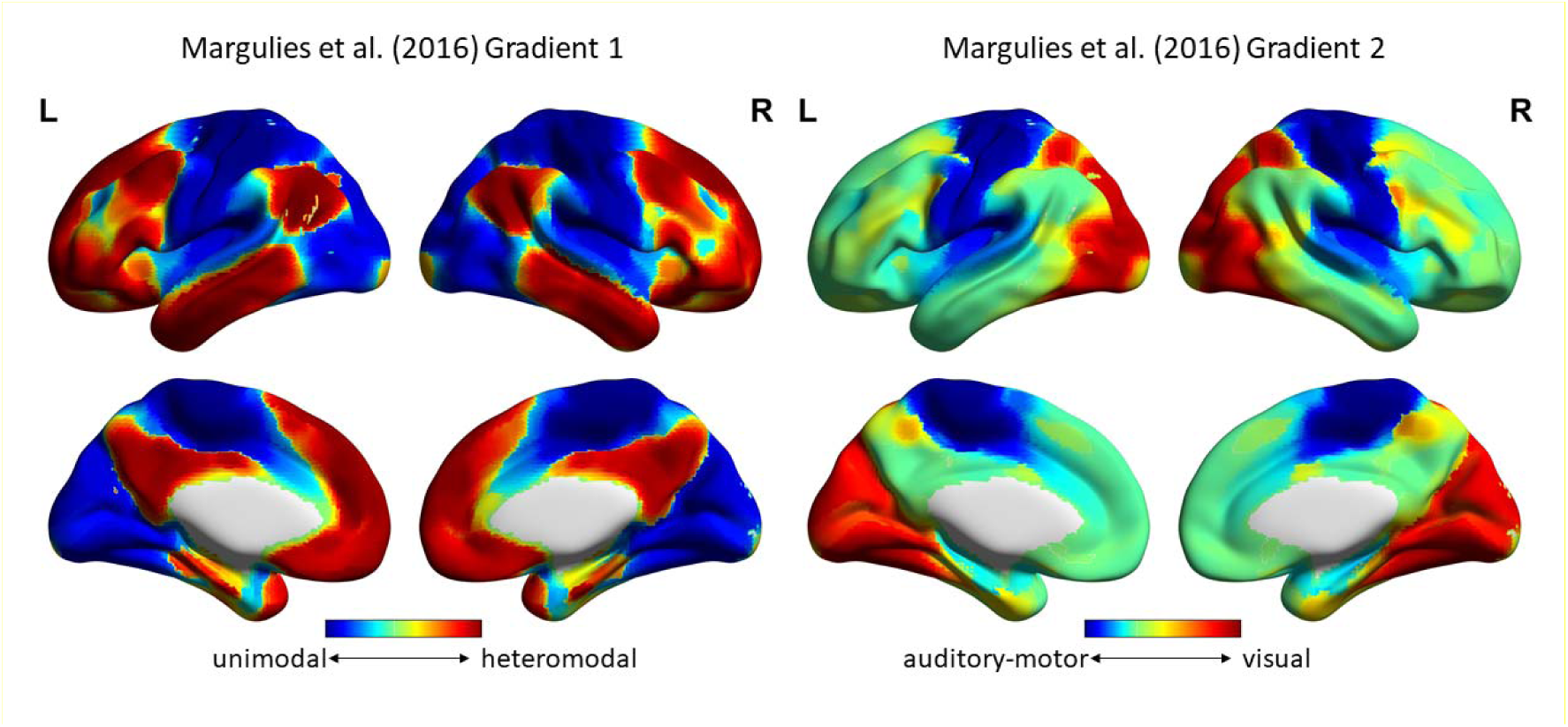
Maps of the Margulies et al. (2016) whole-brain gradients one (G1) and two (G2). The hierarchy of G1 is reflected by the continuous scale from heteromodal regions in red to unimodal regions in blue; for G2, which captures the differentiation of sensorimotor cortices, the scale goes from visual in red to auditory-motor in blue.

In the current research, we examined responses along these two functional gradients within the temporal lobe, re-analysing datasets acquired with MBME fMRI to maximise the signal in this region. These datasets contrasted different kinds of semantic content, allowing us to interrogate the functional semantic responses across the temporal lobe. We used gradients G1 and G2 as defined by Margulies et al. (2016) and masked them by the temporal lobe, assigning voxels to ten decile bins according to values on each gradient, based on intrinsic functional connectivity. The first study examined visually presented verbal semantic judgements and asked participants to link words on the basis of either global semantic relationships (e.g. “seaweed – jellyfish”) or specific visual features (e.g. “meatball – moon”). The second study required participants to generate either emotional states (e.g. *fear* in response to *skydiving*) or meaningful contexts (e.g. *buying something for dinner* in response to *supermarket*) from photographs. Together, the two studies allowed us to investigate whether functional profile varies in a graded fashion across the temporal lobe, or whether there are sharp divisions, in tasks that require participants to make decisions about (written) verbal associations presented to them or actively generate associations to pictures. We examined the effect of semantic content (i.e., association, feature, emotion, context), asking how function varies along the temporal lobe according to the two principal intrinsic connectivity gradients in the functional connectivity of the brain, which capture aspects of processing relevant to temporal cortex and semantic cognition, namely, (i) the continuum from sensorimotor to heteromodal cortex and (ii) the distance from auditory versus visual systems.

The graded hub account predicts that responses along G1 should become more homogeneous moving towards the heteromodal apex of this gradient, while responses further along the gradient may reflect proximity to spoke regions. In contrast, theories advocating a patchwork of function might predict sharp transitions between tasks reflecting specialised function and no homogeneity at the heteromodal end. We can also assess whether activation to verbal semantic stimuli is stronger in regions closer to the auditory end of G2 (despite being presented in the visual domain, given the contribution of auditory-motor processes to language) and whether responses to picture semantic stimuli are stronger at the visual end of G2 (i.e., despite stimulus presentation in the visual domain across both studies, G2 may separate responses according to verbal/picture modality). Furthermore, a recent study found that feature judgments leveraged visual spoke regions more than associative judgments (Chiou, Humphreys, Jung, & Lambon Ralph, 2018); therefore G2 may separate associative and feature judgments along an auditory-visual axis, and on G1 along the unimodal (i.e, feature selection due to access to visual features) to heteromodal (i.e., associative due to more abstract memory codes) processing axis. In addition, generating a semantic context more than an emotion might sit closer to the visual end of G2, given the visuospatial nature of concrete contexts (e.g., contextualising a location such as a supermarket). According to graded theories, both emotion and context generation should rely on heteromodal cortices (e.g., closer to the heteromodal end of G1), as semantic codes are accessed. However, if emotion is more heavily grounded in sensorimotor codes (e.g., Martin, 2016; Niedenthal, Winkielman, Mondillon, & Vermeulen, 2009; Vigliocco et al., 2014), we might see a stronger response on the unimodal end of G1.

## 2. Method

### 2.1. Participants

Participants in Studies 1 and 2 were right-handed, between the ages of 18 and 35, with normal or corrected to normal vision, no history of neurological disorder, and no current psychiatric disorder. Participants were students at the University of York, recruited through word of mouth and participant pools, and paid for their time or awarded course credit. Ethical approval for both studies was granted by the York Neuroimaging Centre at the University of York. Informed consent was obtained from all participants prior to participation. One participant took part in both studies, but there was no further overlap of participants across the two studies.

Study 1 (association/feature judgments): Thirty-four adults were scanned; one participant withdrew from the study due to back pain, another participant was withdrawn due to a structural anomaly, and a further participant was unusable due to the participant falling asleep in both scanning sessions. Two further participants were excluded from data analysis due to poor behavioural performance (2SD’s below the group mean on the feature matching task). Therefore, the final sample consisted of 29 participants (24 female; mean age = 21.1, SD = 3.1). Another 30 native English speakers, who did not take part in the main fMRI experiment, rated the colour and shape similarity and semantic association strength for each word pair (21 females; age range: 18 – 24 years).

Study 2 (emotion/semantic generation): Thirty-three participants were scanned; with one dataset excluded due to the participant not making associations for the majority (52.1%) of trials. The final sample consisted of 32 participants (24 female; mean age = 20.1, SD = 2.4).

### 2.2. Tasks and Paradigms

#### 2.2.1. Study 1: Association and Feature Judgments

In the feature matching task, participants made yes/no decisions about whether two visually presented words shared a particular visual feature (colour or shape; Figure 2). The trials were created such that participants would respond yes to roughly half of the trials, allowing us to separate the neural response to yes and no decisions. For example, for colour matching: participants would respond ‘yes’ to “DALMATIAN – COW”, due to their colour similarity, but ‘no’ to “COAL -TOOTH” due to the lack of colour feature similarity. The task was split into four runs, with two feature conditions (colour/shape), presented in a mixed design. The feature type was split into four mini blocks (2mins 30s each), resulting in a total run time of 10.55mins. In each mini block, 20 trials were presented in a rapid event-related design. In order to maximize the statistical power, the stimuli were presented with a temporal jitter randomized from trial to trial (Dale, 1999), with a variable inter-trial of 3 to 5 s. Each trial started with a fixation, followed by the feature type (colour/shape) at the top of the screen, and the concepts presented centrally (Figure 2, detailed task schematic in supplementary materials Figure S1). These remained on-screen until the participant responded, or for a maximum of 3 s. The condition order was counterbalanced across runs and run order was counterbalanced across participants.

**Figure 2.**
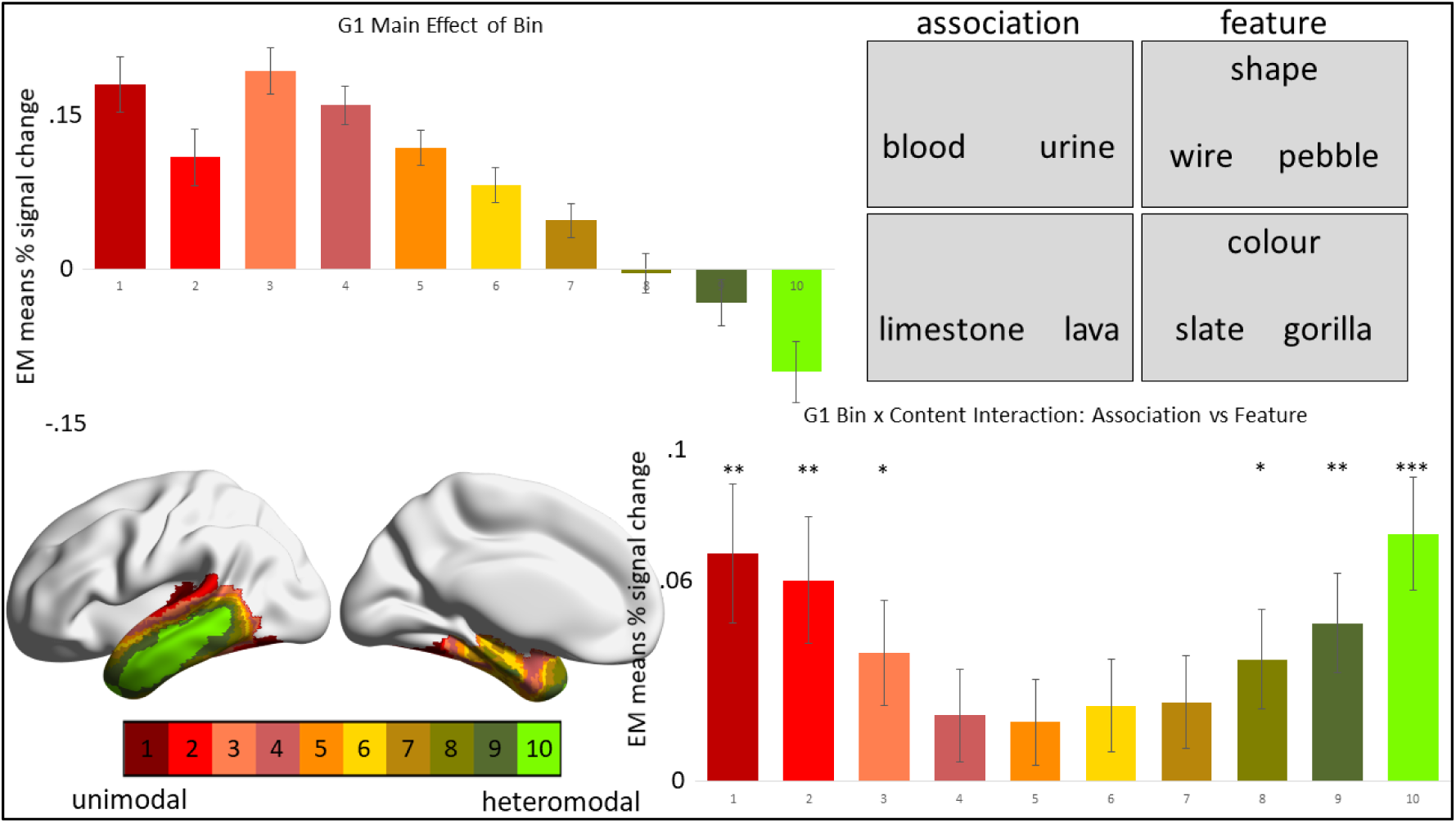
Results for G1 constrained to left temporal lobe for associative and feature judgments. Top left: Main effect of bin for associative and feature judgments, the y-axis represents estimated marginal means of percent signal change for each decile bin (x-axis). Top right: Task example for associative and feature judgments. Bottom left: Left temporal lobe gradient 1 segmented into 10 decile bins. Bottom right: The interaction of bin and content for associative and feature judgments – the bars represent the difference between associative – feature activation in each bin (x-axis) based on estimated marginal means of percent signal change (y-axis). The gradient bin colour scale is the same across brain and graphical representations. \**p* < .05, \*\**p* < .01, \*\*\**p* ≤ .001, for post-hoc Bonferroni t-tests, which can also be found in Table 1.

**Table 1.** Bonferroni-corrected post-hoc t-tests for L temporal lobe G1 and G2.

|  | bin | EM Means |  | Mean Difference<br>(Association-Feature) | Std.<br>Error | Sig.<sup>b</sup> | 95% Confidence<br>Interval for<br>Difference<sup>b</sup> |  |
| --- | --- | --- | --- | --- | --- | --- | --- | --- |
|  |  | Assoc | Feat |  |  |  | Lower<br>Bound | Upper<br>Bound |
| G1 | **1** | 0.212 | 0.144 | **.069\*** | **0.021** | **0.003** | **0.025** | **0.112** |
|  | **2** | 0.138 | 0.077 | **.061\*** | **0.019** | **0.003** | **0.022** | **0.099** |
|  | **3** | 0.21 | 0.172 | **.039\*** | **0.016** | **0.022** | **0.006** | **0.071** |
|  | 4 | 0.168 | 0.148 | 0.02 | 0.014 | 0.158 | -0.008 | 0.048 |
|  | 5 | 0.126 | 0.107 | 0.018 | 0.013 | 0.178 | -0.009 | 0.045 |
|  | 6 | 0.093 | 0.07 | 0.023 | 0.014 | 0.108 | -0.005 | 0.05 |
|  | 7 | 0.059 | 0.035 | 0.024 | 0.014 | 0.089 | -0.004 | 0.052 |
|  | **8** | 0.014 | -0.023 | **.037\*** | **0.015** | **0.019** | **0.006** | **0.067** |
|  | **9** | -0.008 | -0.055 | **.048\*** | **0.015** | **0.003** | **0.018** | **0.078** |
|  | **10** | -0.061 | -0.136 | **.075\*** | **0.017** | **<.001** | **0.041** | **0.109** |
| G2 | **1** | 0.013 | -0.045 | **.058\*** | **0.022** | **0.012** | **0.014** | **0.103** |
|  | **2** | 0.033 | -0.015 | **.049\*** | **0.017** | **0.008** | **0.014** | **0.084** |
|  | **3** | 0.02 | -0.031 | **.051\*** | **0.014** | **0.001** | **0.021** | **0.08** |
|  | **4** | -0.018 | -0.074 | **.056\*** | **0.015** | **0.001** | **0.025** | **0.088** |
|  | **5** | -0.034 | -0.093 | **.060\*** | **0.015** | **0.001** | **0.028** | **0.091** |
|  | **6** | 0.002 | -0.048 | **.050\*** | **0.015** | **0.002** | **0.02** | **0.081** |
|  | 7 | 0.07 | 0.049 | 0.02 | 0.014 | 0.167 | -0.009 | 0.049 |
|  | 8 | 0.121 | 0.108 | 0.012 | 0.013 | 0.336 | -0.013 | 0.038 |
|  | 9 | 0.241 | 0.242 | -0.001 | 0.015 | 0.972 | -0.031 | 0.03 |
|  | **10** | 0.503 | 0.447 | **.056\*** | **0.026** | **0.042** | **0.002** | **0.11** |

Based on estimated marginal means; \* The mean difference is significant at the .05 level;
<sup>b</sup>adjustment for multiple comparisons: Bonferroni.

In the association task, participants made yes/no decisions about whether two visually presented words shared a semantic association or not. The trials were created such that participants would respond yes to roughly half of the trials, allowing us to separate the neural response to yes and no decisions. The same stimuli were used across both semantic feature matching and semantic associations task. For example, “DALMATIAN – COW’ are semantically related (and also have similar colour) whereas “COAL – PUMA” are not semantically related (but still share the colour feature). This task was split into four runs, presented in a rapid event-related design. Each run consisted of 80 trials (10.35 mins per run), and the procedure was the same as the feature matching task except only two words were presented on the screen (as no condition cue was needed).

Half of the participants responded with their right index finger to indicate yes and with their middle finger to indicate no; the other half pressed the opposite buttons. The feature and association task sessions were separated by one week, and their order was counterbalanced across participants.

#### 2.2.2. Study 2: Emotion and Context Generation and Switch

This study required participants to generate context or emotion associations to pictures, and then retrieve a new association to the same picture in a second phase. Stimuli were taken from the International Affective Picture System (IAPS; Lang, Brady, & Cuthbert, 2008), a database of pictures normed for valence and arousal. Thirty-six pictures were selected for *emotion* associations and were either positive (valence mean > 6) or negative (valence mean < 4), with an equal number of positive and negative images in each experimental run. A further thirty-six pictures were selected for *context* associations, all of neutral valence (mean valence between 4 and 6). Valence for pictures across conditions was significantly different (negative emotion < *context* < positive *emotion* pictures; U < 1, *p* < .001). Ratings of mean arousal were significantly higher for *emotion* than *context* pictures (U = 221.0, *p* < .001), but were matched between positive and negative *emotion* images (U = 153.5, *p* = .791). Identifiers for each image and normed ratings of valence and arousal are provided in Supplementary Table S1; these data have been used in a previous study focussing on default mode network subdivisions (Souter et al., 2023).

Trials were presented in mini-blocks of association type (emotion/context) with six trials in each mini-block per run. The order of trials within a run was consistent across participants, but run order was counterbalanced across participants. Each mini-block started with a 2 second instruction (i.e., ‘SEMANTIC’ or ‘EMOTION’), and each trial started with a jittered fixation cross (1-3 s). During the ‘GENERATE’ phase participants were presented with a picture; during *context* associations, participants identified a meaningful context from their general knowledge (i.e., they were asked not to rely on a specific episodic memory), and for emotion associations, participants were asked to embody emotions evoked by the image when generating the association. The ‘SWITCH’ phase required participants to stop reflecting on their initial association, and generate a new one. The same image was used in both generate and switch phases of the trial. Trial time across g*enerate* and *switch* phases was jittered between 3.5 and 6.5s, with an average of 5s. After each *generate* and *switch* phase, participants rated the strength of their self-generated association on a scale from 1 (no real relationship) to 7 (very strong relationship). A final rating of switch difficulty (i.e., how difficult it was to switch from initial association to a new one) was given at the end of the whole trial on a scale from 1 (very easy) to 7 (very difficult). Each rating period lasted 3s (Figure 4; detailed task schematic in supplementary materials Figure S1).

**Table 2.**
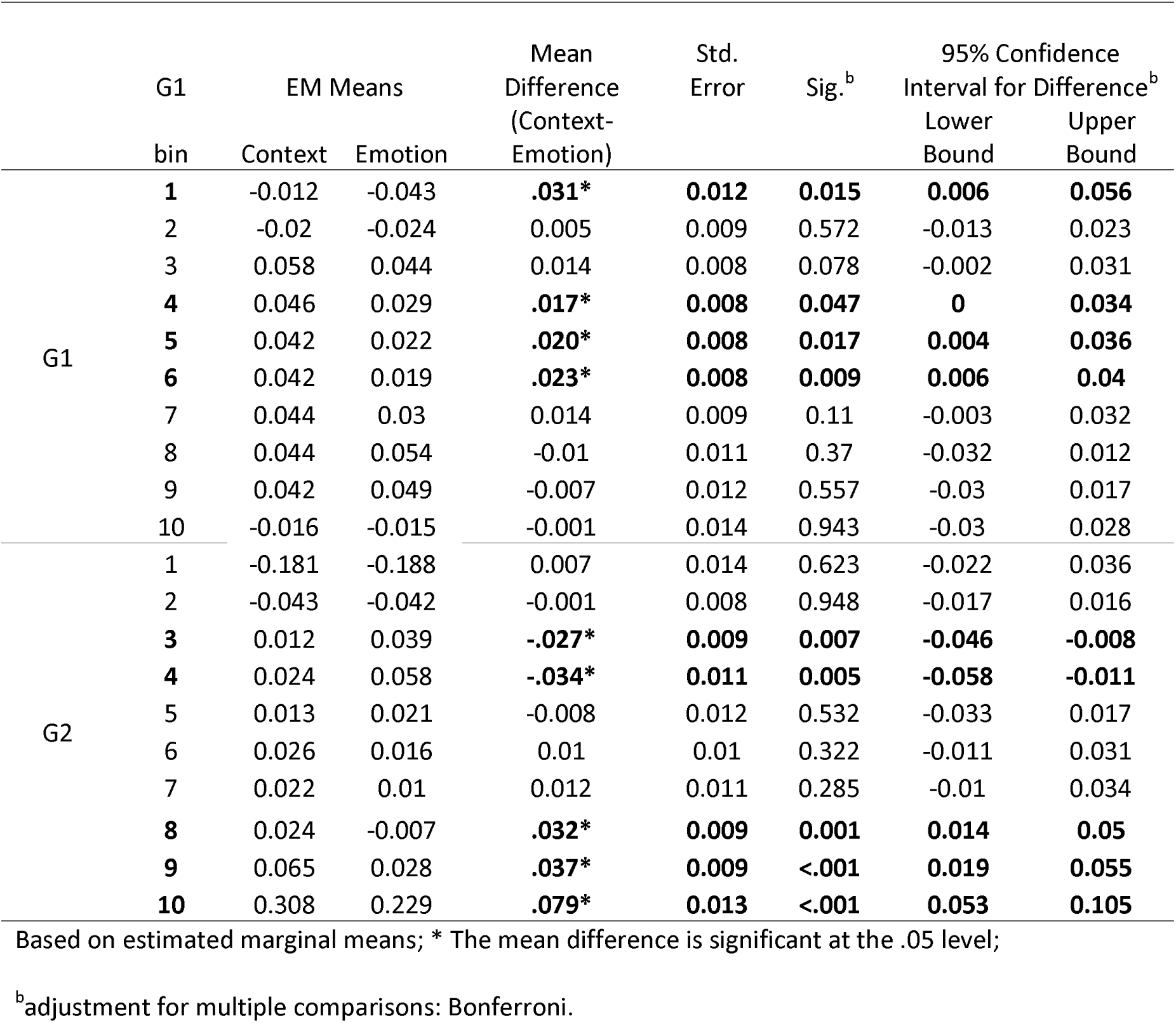
Bonferroni-corrected post-hoc t-tests for L temporal lobe G1 and G2.

|  | G1 | EM Means |  | Mean Difference (Context-Emotion) | Std. Error | Sig.<sup>b</sup> | 95% Confidence Interval for Difference<sup>b</sup> |  |
| --- | --- | --- | --- | --- | --- | --- | --- | --- |
|  | bin | Context | Emotion |  |  |  | Lower Bound | Upper Bound |
| G1 | 1 | -0.012 | -0.043 | **.031\*** | **0.012** | **0.015** | **0.006** | **0.056** |
|  | 2 | -0.02 | -0.024 | 0.005 | 0.009 | 0.572 | -0.013 | 0.023 |
|  | 3 | 0.058 | 0.044 | 0.014 | 0.008 | 0.078 | -0.002 | 0.031 |
|  | 4 | 0.046 | 0.029 | **.017\*** | **0.008** | **0.047** | **0** | **0.034** |
|  | 5 | 0.042 | 0.022 | **.020\*** | **0.008** | **0.017** | **0.004** | **0.036** |
|  | 6 | 0.042 | 0.019 | **.023\*** | **0.008** | **0.009** | **0.006** | **0.04** |
|  | 7 | 0.044 | 0.03 | 0.014 | 0.009 | 0.11 | -0.003 | 0.032 |
|  | 8 | 0.044 | 0.054 | -0.01 | 0.011 | 0.37 | -0.032 | 0.012 |
|  | 9 | 0.042 | 0.049 | -0.007 | 0.012 | 0.557 | -0.03 | 0.017 |
|  | 10 | -0.016 | -0.015 | -0.001 | 0.014 | 0.943 | -0.03 | 0.028 |
| G2 | 1 | -0.181 | -0.188 | 0.007 | 0.014 | 0.623 | -0.022 | 0.036 |
|  | 2 | -0.043 | -0.042 | -0.001 | 0.008 | 0.948 | -0.017 | 0.016 |
|  | 3 | 0.012 | 0.039 | **-.027\*** | **0.009** | **0.007** | **-0.046** | **-0.008** |
|  | 4 | 0.024 | 0.058 | **-.034\*** | **0.011** | **0.005** | **-0.058** | **-0.011** |
|  | 5 | 0.013 | 0.021 | -0.008 | 0.012 | 0.532 | -0.033 | 0.017 |
|  | 6 | 0.026 | 0.016 | 0.01 | 0.01 | 0.322 | -0.011 | 0.031 |
|  | 7 | 0.022 | 0.01 | 0.012 | 0.011 | 0.285 | -0.01 | 0.034 |
|  | 8 | 0.024 | -0.007 | **.032\*** | **0.009** | **0.001** | **0.014** | **0.05** |
|  | 9 | 0.065 | 0.028 | **.037\*** | **0.009** | **<.001** | **0.019** | **0.055** |
|  | 10 | 0.308 | 0.229 | **.079\*** | **0.013** | **<.001** | **0.053** | **0.105** |

Based on estimated marginal means; \* The mean difference is significant at the .05 level;
<sup>b</sup>adjustment for multiple comparisons: Bonferroni.

Immediately following the scan, participants completed a ‘recall’ assessment. Pictures were presented in the same order as in the scanner, and participants typed the contexts or emotions they generated, as well as rating confidence in recall (from 1-7), for both the *generate* and *switch* phases. This data was used to qualitatively validate that participants performed the task as intended.

For both Studies 1 and 2, participants completed practice sessions the day before their scanning session via Zoom. Python scripts used for presentation of the semantic tasks are available on Open Science Framework (OSF; Study 1 at https://osf.io/p2s3w/ and Study 2 at https://osf.io/498ur/).

### 2.3 Temporal Lobe Gradients

Whole-brain decompositions of intrinsic connectivity, termed ‘gradients’, capture dimensions of intrinsic connectivity change across the cortical surface. We masked Margulies et al.’s (2016) gradients 1 (G1) and 2 (G2) by the temporal lobe, derived from the MNI-maxprob-thr50-2mm mask, to capture these intrinsic connectivity gradients in the temporal lobe. First, the temporal lobe mask was split into left and right and multiplied by the Margulies et al. (2016) G1 and G2 (generating 4 temporal lobe gradients: Left G1, Right G1, Left G2, Right G2). Next, the temporal lobe gradients were divided into ten decile bins, according to their values on the connectivity gradient. Then, percent signal change was calculated for each bin in each participant (and condition) and subjected to within-subjects ANOVAs (all interactions with bin were Greenhouse-Geisser corrected). While this study focused on the gradients within the temporal lobe, the results for whole brain gradient ANOVAs (G1 and G2) can be found in supplemental materials (Table S2 and S3).

### 2.4. Image Acquisition

Whole brain structural and functional MRI data were acquired using a 3T Siemens MRI scanner utilising a 64-channel head coil, tuned to 123 MHz at the York Neuroimaging Centre, University of York. We used a multiband-multiecho (MBME) EPI sequence with the following parameters: TR=1.5 s; TEs = 12, 24.83, 37.66 ms; 48 interleaved slices per volume with slice thickness of 3 mm (no slice gap); FoV = 24 cm (resolution matrix = 3x3x3; 80x80); 75° flip angle; volumes per run below; 7/8 partial Fourier encoding and GRAPPA (acceleration factor = 3, 36 ref. lines; multi-band acceleration factor = 2). Structural T1-weighted images were acquired using an MPRAGE sequence (TR = 2.3 s, TE = 2.26 s; voxel size = 1x1x1 isotropic; 176 slices; flip angle = 8°; FoV= 256 mm; interleaved slice ordering). The number of runs, run time and volumes collected per run are as follows: feature task = 4 runs x 10.55 mins (422 volumes); association task = 4 runs x 10.35 (414 volumes); emotion/context task = 6 runs x 4.5 mins (180 volumes).

For the association/feature study, we also collected a high-resolution T2-weighted (T2w) scan using an echo-planar imaging sequence (TR = 3.2 s, TE = 56 ms, flip angle = 120°; 176 slices, voxel size = 1x1x1 isotropic; Fov = 256 mm), and a 9 minute resting-state scan (acquired using an EPI sequence; 80° flip angle; GRAPPA acceleration factor of 2; resolution matrix = 3x3x4; 64x64; TR = 3s, TE = 15 ms, FoV = 192 mm) that were not used in the current project.

### 2.5. Image Pre-processing

A MBME sequence was used to optimise signal from the anterior and medial temporal regions, while maintaining optimal signal across the whole brain (Halai et al., 2014). We used TE Dependent ANAlysis (tedana; version 0.0.12; Kundu et al., 2013; The tedana Community et al., 2021; Kundu, Inati Sj Fau - Evans, Evans Jw Fau - Luh, Luh Wm Fau -Bandettini, & Bandettini, 2012) to combine the images. Anatomical pre-processing (fsl_anat; https://fsl.fmrib.ox.ac.uk/fsl/fslwiki/fsl_anat) included re-orientation to standard MNI space (fslreorient2std), automatic cropping (robustfov), bias-field correction (RF/B1 – inhomogeneity-correction, using FAST), linear and non-linear registration to standard-space (using FLIRT and FNIRT), brain extraction (using FNIRT, BET), tissue-type segmentation (using FAST) and subcortical structure segmentation (FAST). The multi-echo data were pre-processed using AFNI (https://afni.nimh.nih.gov/), including de-spiking (3dDespike), slice timing correction (3dTshift; heptic interpolation), and motion correction (3dvolreg applied to echo 1 to realign all images to the first volume; these transformation parameters were then applied to echoes 2 and 3; cubic interpolation). Runs with motion greater than 1.1mm (absolute) were excluded from analyses across both semantic studies. This resulted in the removal of the final run for two participants for the feature matching task and for one participant for the association task. Relative displacement was less than .18mm across both Studies 1 and 2.

### 2.7. fMRI Data Analysis

Individual level analyses were conducted using FSL-FEAT version 6 (FMRIB’s Software Library, www.fmrib.ox.ac.uk/fsl; Jenkinson, Bannister, Brady, & Smith, 2002; Smith et al., 2004; Woolrich et al., 2009). Denoised optimally-combined time series output from tedana were submitted to FSL and pre-processing included high-pass temporal filtering (Gaussian-weighted least-squares straight line fitting, with sigma = 50s), linear co-registration to the corresponding T1-weighted image and to MNI152 standard space (Jenkinson & Smith, 2001), spatial smoothing using a Gaussian kernel with full-width-half-maximum of 6 mm, and grand-mean intensity normalisation of the entire 4D dataset by a single multiplicative factor. Pre-processed time series data were modelled using a general linear model correcting for local autocorrelation (Woolrich, Ripley, Brady, & Smith, 2001). No motion parameters were included in the models, as the data had already been denoised as part of the TEDANA pipeline (Kundu et al., 2012).

Study 1 examined association and feature judgments. EVs were as follows: (1) the mean activity to yes responses, (2) mean for no responses, (3) parametric effects for yes responses, (4) parametric effects for no responses. Parametric regressors were derived from an independent set of participants (see Participants). For the association task, we included rated association strength for each word pair on a five-point scale (1 = not at all related to 5 = very related). For the feature matching task, we examined rated feature similarity of the pair of items (1 = not similar at all to 5 = very similar). These parametric EVs are not relevant to the current study. A fifth EV captured any incorrect (feature matching) or missed (association/feature) trials. Response type (yes/no) was entered into all ANOVAs for Study 1, however this variable was beyond the scope of this study and did not consistently interact with gradient bins across G1 or G2 (see Supplementary Materials, Table S4).

Study 2 examined emotion and context across generation and switch phases. EVs were as follows: (1) context generate, (2) context switch, (3) emotion generate, (4) emotion switch, (5) all self-report rating periods and (6) condition prompt from the start of each mini-block, (7) parametric context switch difficulty, (8) parametric emotion switch difficulty. These parametric EVs are not relevant to the current study. The generate and switch conditions were averaged within content type (i.e., emotion, context) to create a COPE (contrast of parameter estimates) for emotion and context trials.

Whole brain results for these studies are reported elsewhere, please see Wang et al. (2023) for Study 1 and Souter et al. (2023) for Study 2. We extracted percent signal change for each gradient bin within the temporal lobe in Studies 1 and 2.

## 3. Results

### 3.1.1. G1: Association versus Feature Matching

An ANOVA of the 10 bins (of G1 in left temporal lobe) across two tasks (association/feature) and decision type (yes/no) revealed a significant main effect of bin (F(2.8, 79) = 61.7 *p* < .001, η_p_ ^2^ = .69; Figure 2), with a linear trend (F1, 28) = 107.2, *p* < .001, η_p_ ^2^ = .79), indicating a gradual shift in activation from more unimodal parts of temporal cortex to deactivation in heteromodal temporal cortex. This analysis also revealed an interaction of bin by content (F(2.5, 71) = 5.2 *p* = .004, η_p_ ^2^ = .16; Figure 2), with a quadratic trend (F1, 28) = 19.4, *p* < .001, η_p_ ^2^ = .41), reflecting u-shaped differential activation between association and feature judgments along G1. Full ANOVA tables can be found in supplementary materials (Table S4; Table S6 for trends).

Post-hoc Bonferroni corrected t-tests confirmed differences between the two content types, with significantly more activation for the association decisions in bins 1, 2 and 3 (the more unimodal end of the temporal G1) and more deactivation of feature semantic tasks in the heteromodal end of left temporal cortex (Table 1; Figure 2).

### 3.1.2. G2: Association versus Feature Matching

An ANOVA of the 10 bins (of G2 in left temporal lobe) by task type (association/feature) by decision type (yes/no) revealed a significant main effect of bin (F(3.3, 93) = 142.9 *p* < .001, η_p_ ^2^ = .84; Figure 3), with a linear trend (F1, 28) = 291.6, *p* < .001, η_p_ ^2^ = .91) reflecting a shift from bin 5 (deactivation) to 10 (visual end; activation). This analysis also revealed an interaction of bin by task (F(2.4, 67) = 3.6 *p* = .024, η_p_ ^2^ = .12; Figure 3), with a cubic trend reflecting differences in deactivation towards the auditory end, and a difference in activation at the visual extreme (see post-hoc tests reported below), with no difference in between. Full ANOVA tables can be found in the supplementary materials (Table S4; Table S6 for trends).

**Figure 3.**
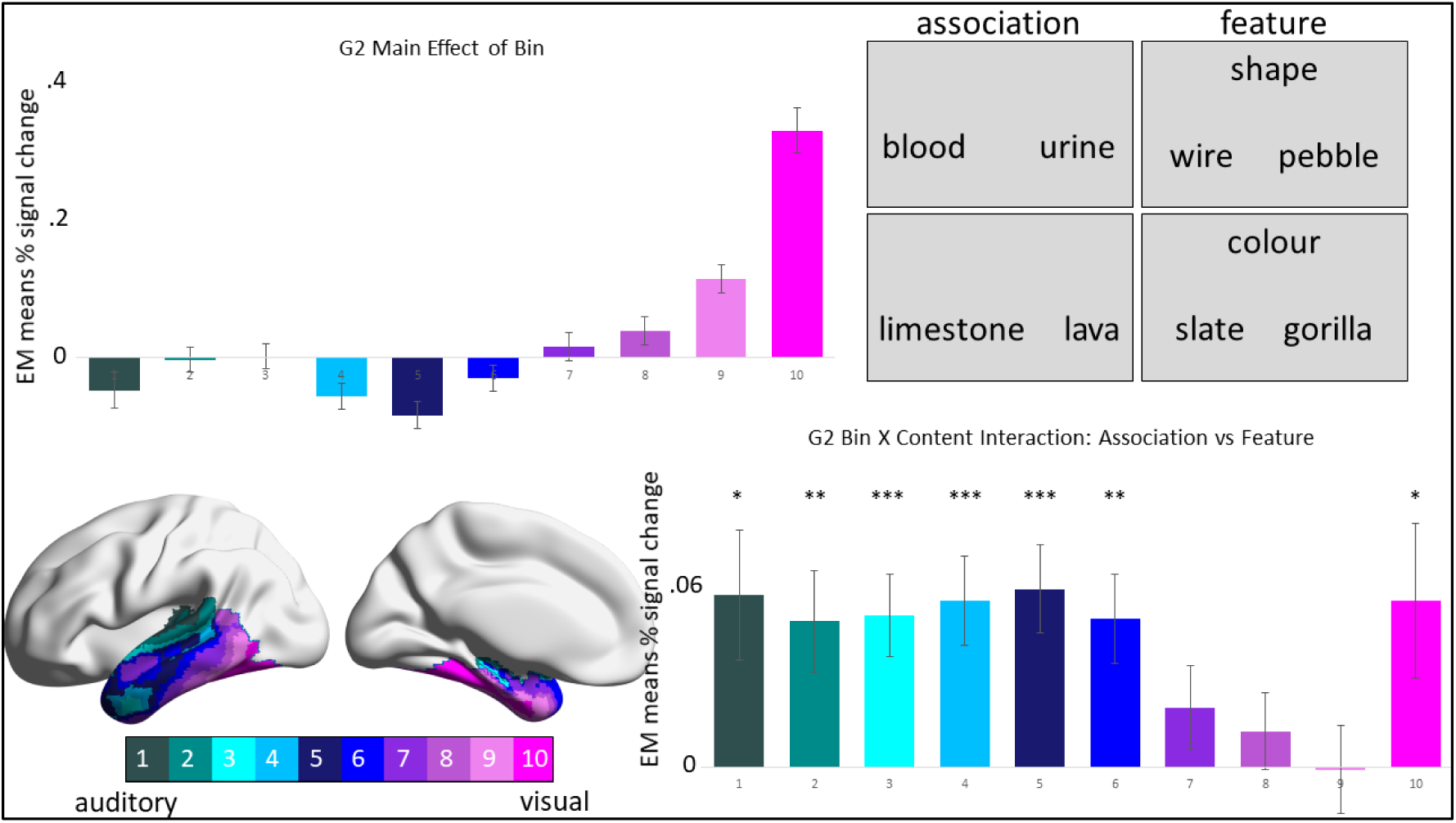
Results for G2 constrained to left temporal lobe for associative and feature judgments. Top left: Main effect of bin for associative and feature judgments, the y-axis represents estimated marginal means of percent signal change for each decile bin (x-axis). Top right: Task example for associative and feature judgments. Bottom left: Left temporal lobe gradient 2 segmented into 10 decile bins. Bottom right: The interaction of bin by content for associative and feature judgments – the bars represent the difference between associative – feature activation in each bin (x-axis) based on estimated marginal means of percent signal change (y-axis). The gradient bin colour scale is the same across brain and graphical representations. \**p* < .05, \*\**p* < .01, \*\*\**p* ≤ .001, for post-hoc Bonferroni t-tests, which can also be found in Table 1.

Post-hoc Bonferroni corrected t-tests confirmed differences between the two tasks. Bins 1, 3, 4, (towards the auditory end of G2) and 6 showed more deactivation for feature semantic decisions. In bin 2 there was a significant difference between the two tasks, with activation for the association judgments, but not feature matching. In bin 5, both tasks elicited deactivation, but with significantly more deactivation for feature than association judgments. Both tasks activated bins 7, 8, and 9, with no significant differences; however, bin 10 at the visual extreme of the gradient showed significantly more activation for association than feature judgments (Table 1; Figure 3).

#### 3.2.1. G1: Context versus Emotion Generation

An ANOVA of the 10 bins (of G1 in left temporal lobe) by content type (context/emotion) revealed a significant main effect of bin (F(2.1, 66) = 5.3 *p* = .006, η_p_ ^2^ = .15; Figure 4), with a cubic trend (F(1, 31) = 24.1, *p* < .001, η_p_ ^2^ = .44) reflecting deactivation at the two extreme ends of temporal lobe G1 (in bins unimodal 1 and 2; and heteromodal 10) and a u-shaped activation profile from bins 3 to 9. This analysis also revealed an interaction of bin by content (F(2.4, 76) = 4.5 *p* = .01, η_p_ ^2^ = .13; Figure 4), with a weak and non-significant linear trend (F(1, 31) = 3.8, *p* = .061, η_p_ ^2^ = .11). Full ANOVA tables can be found in supplementary materials (Table S5; Table S7 for trends).

Post-hoc Bonferroni corrected t-tests confirmed differences between the two content types, with significantly less deactivation for context than emotion decisions at the unimodal end (bin 1) and significantly more activation for context generation in the middle of the gradient (bins 4, 5, 6; Table 2; Figure 4).

**Figure 4.**
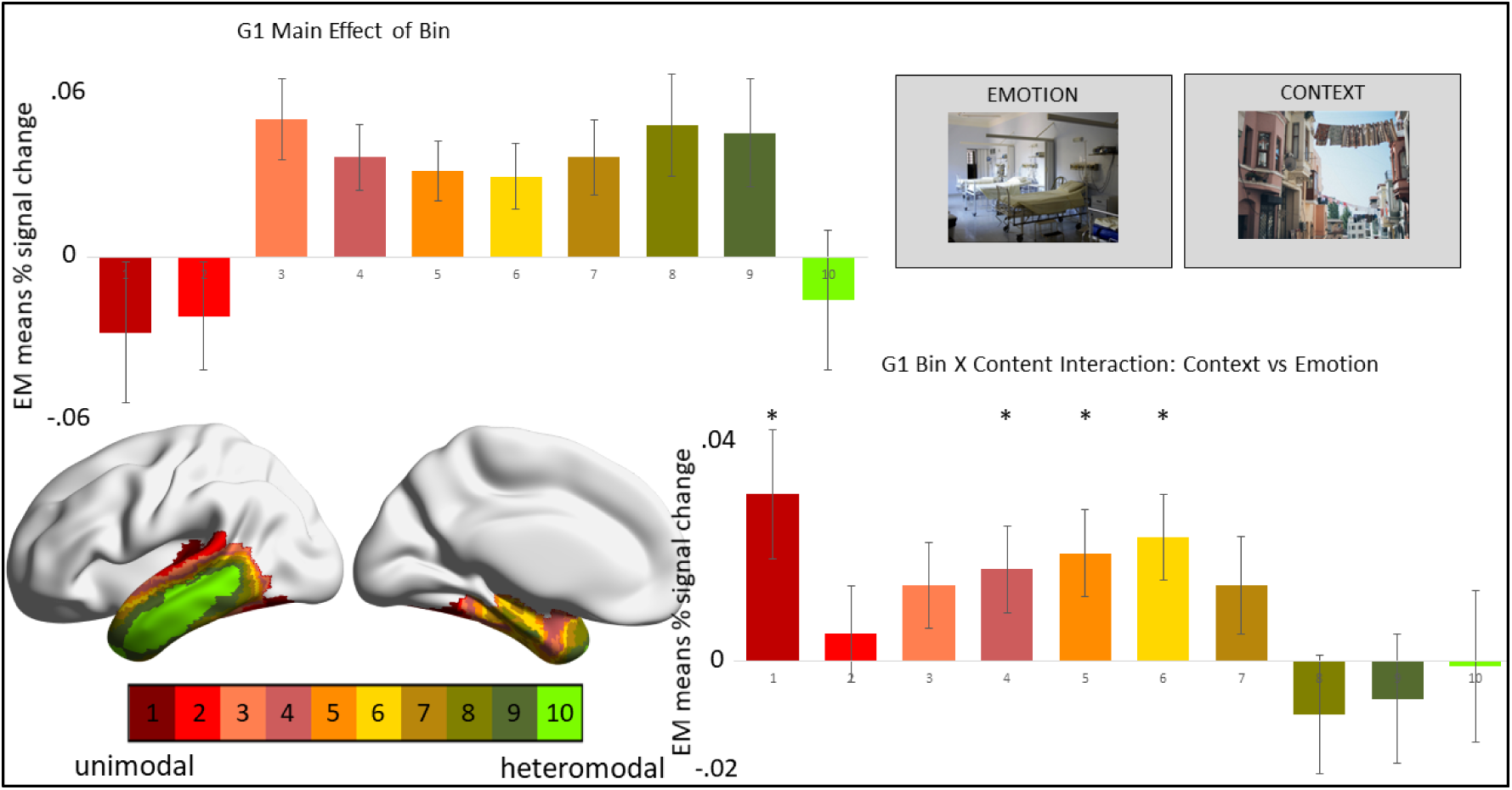
Results for G1 constrained to left temporal lobe for context and emotion generation. Top left: Main effect of bin for context and emotion generation, the y-axis represents estimated marginal means of percent signal change for each decile bin (x-axis). Top right: Task example for context and emotion generation. Bottom left: Left temporal lobe gradient 2 segmented into 10 decile bins. Bottom right: The interaction of bin by content for context and emotion generation – the bars represent the difference between context – emotion activation in each bin (x-axis) based on estimated marginal means of percent signal change (y-axis). The gradient bin colour scale is the same across brain and graphical representations. \**p* < .05, \*\**p* < .01, \*\*\**p* ≤ .001, for post-hoc Bonferroni t-tests, which can also be found in Table 2.

#### 3.2.2. G2: Context versus Emotion Content Generation

An ANOVA of the 10 bins (of G2 in left temporal lobe) by content type (context/emotion) revealed a significant main effect of bin (F(2.8, 87) = 65.9 *p* < .001, η_p_ ^2^ = .68; Figure 5), with linear (F(1, 31) = 163 *p* < .001, η_p_ ^2^ = . 48) and quadratic trends (F(1, 31) = 155.5 *p* < .001, η_p_ ^2^ = .83) sharing similar F-values, reflecting a sharp change from deactivation at the auditory end of temporal G2 to activation at the visual end, with relatively little change between bins 2-9. This analysis also revealed an interaction of bin by content (F(3.8, 119) = 18.2 *p* < .001, η_p_^2^ = .37; Figure 5), which was linear (F(1, 31) = 45.7 *p* < .001, η_p_ ^2^ = .6). Full ANOVA tables can be found in supplementary materials (Table S5; Table S7 for trends).

**Figure 5.**
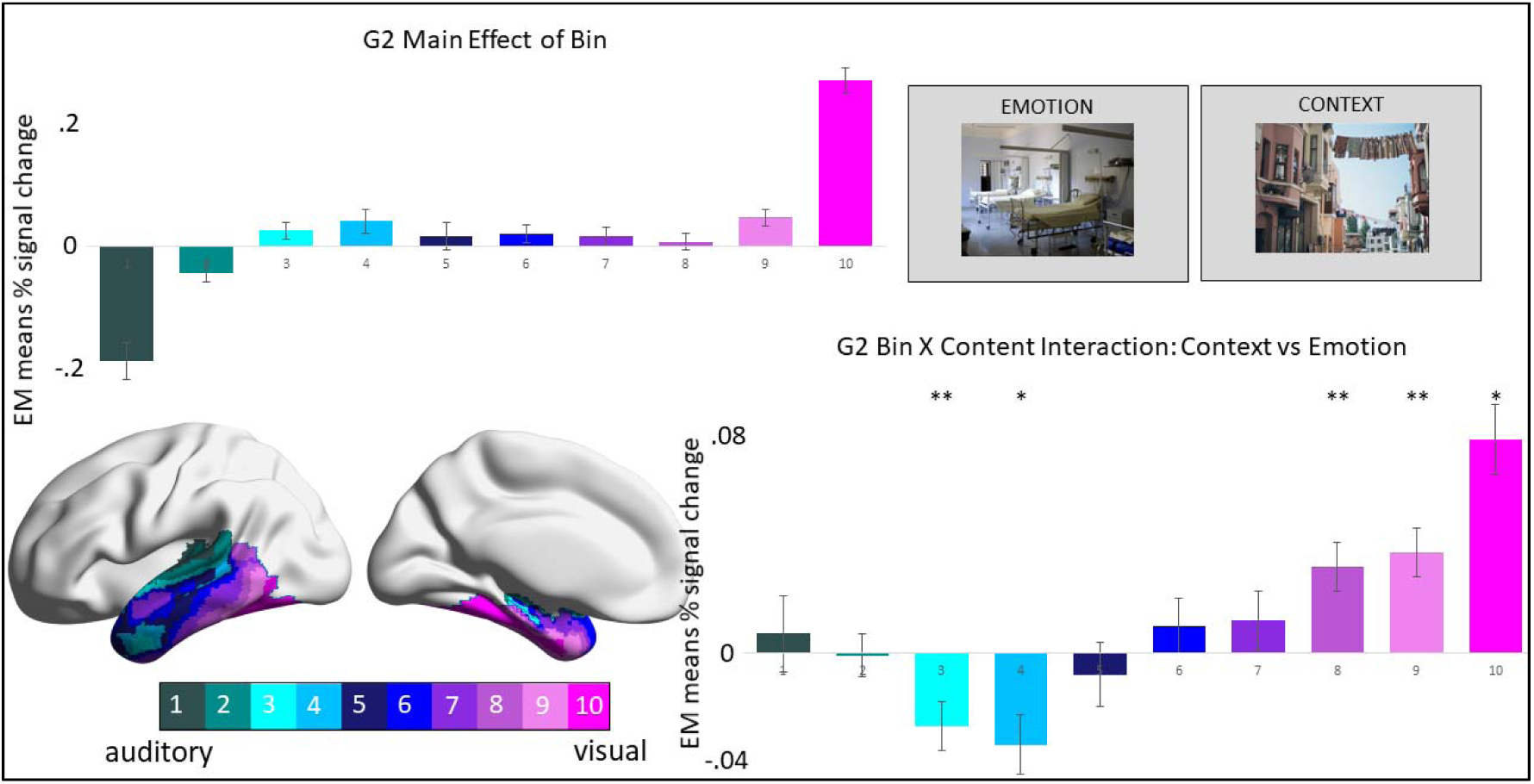
Results for G2 constrained to left temporal lobe for context and emotion generation. Top left: Main effect of bin for context and emotion generation, the y-axis represents estimated marginal means of percent signal change for each decile bin (x-axis). Top right: Task example for context and emotion generation. Bottom left: Left temporal lobe gradient 2 segmented into 10 decile bins. Bottom right: The interaction of bin by content for context and emotion generation – the bars represent the difference between context – emotion activation in each bin (x-axis) based on estimated marginal means of percent signal change (y-axis). The gradient bin colour scale is the same across brain and graphical representations. \**p* < .05, \*\**p* < .01, \*\*\**p* ≤ .001, for post-hoc Bonferroni t-tests, which can also be found in Table 2.

Post-hoc Bonferroni corrected t-tests confirmed differences between the two content types. The emotion task activated bins 3 and 4 towards the auditory end of temporal lobe G2. Both emotion and context semantics activated bins 9 and 10, but context-based semantics activated bins 8, 9 and 10 significantly more than emotion (Table 2; Figure 5), demonstrating greater activation for context-based semantics at the visual end of left temporal lobe G2.

### 3.3 G1 and G2 in Left vs Right Temporal Lobe

We also interrogated G1 and G2 in the right temporal lobe. No substantial differences were found in G1; these data are included as supplementary materials. To summarise, in temporal G1 there was a marginal and non-significant three-way interaction of hemisphere by bin by content (F(3.9, 109.6) = 2.3, *p* = .07, η_p_ ^2^ = .075) for association and feature judgments and a non-significant effect for emotion and context (F(3.1, 97.4) = 1.5, *p* = .23, η_p_ ^2^ = .05). In temporal G2 there was a significant interaction of hemisphere by bin by association/feature matching (F(3.8, 105.3) = 3.8, *p* = .007, η_p_^2^ = .12), reflecting greater activation across bins for the left hemisphere, with the size of this difference increasing closer to the visual end of the gradient; Tables S8; S10). There was also a significant interaction of hemisphere by bin by context/emotion generation (F(4.6, 142.6) = 4.6, *p* = .001, η_p_ ^2^ = .13), reflecting greater activation in middle portions of G2 (bins 3-7) for the left hemisphere, and greater activation in the right hemisphere at the visual end (bins 9 and 10) of this gradient for both context and emotion (Tables S9; S10).

## 4. Discussion

This investigation highlights the complex nature of temporal lobe organisation, with much work still to be done. We found common effects of gradient location within each study, but also subtle differences across gradients dependent on the task condition. Furthermore, we found that differences in the functional response to different types of semantic content were often in the relative *magnitude* of activation/deactivation, rather than a binary on/off pattern dissociable by content. In G1 within temporal lobe we found that, overall, activation decreased for associative and feature judgments to visually presented verbal stimuli in a graded fashion moving away from unimodal regions towards heteromodal cortex. However, within this overall pattern, we also found that associative judgments decreased activation in heteromodal cortex significantly *less* than feature semantic judgments. However, the associative judgments also activated the unimodal end of G1 significantly more than the feature judgments, suggesting a complex pattern leveraging both “spoke” and heteromodal cortex. The activation along the middle of G1 was not significantly different between associative and feature content, changing in a linear fashion along the gradient.

G2 allowed us to further interrogate how the unimodal end of G1 might be recruited for different types of semantic judgment. This analysis again showed a clear main effect of gradient location, such that activation transitioned from auditory deactivation to strong visual activation on G2. The differences in magnitude between associative and feature judgments revealed that associative judgments were significantly less likely to deactivate the auditory end of G2, while also activating the visual end more strongly than the feature judgments. Taken together, the visually presented verbal judgments to both associative and feature content demonstrated a graded functional change along the temporal lobe, according to the intrinsic connectivity profile of this region (as measured by G1 and G2).

However, activation along G1 was not graded for context and emotion generation – this type of internally generated thought to externally presented pictures deactivated the two extreme ends of G1 (bins 1 and 2 on the unimodal end; bin 10 on the heteromodal end), and activated the middle of G1 in a U-shaped fashion. Subtle differences emerged along the gradient between the two content types: emotion content deactivated the unimodal end of G1 (bin 1), while the middle of this gradient was more strongly activated by context than emotion generation. There were no significant differences in activation between context and emotion towards heteromodal cortex, possibly due to the need to generate semantic associations in both conditions (Andrews-Hanna, Smallwood, & Spreng, 2014; Smallwood et al., 2021; Smith, Mitchell, & Duncan, 2018). Again, G2 allowed us to investigate whether activation was graded according to intrinsic connectivity profiles aligned with sensorimotor cortices: like study 1, we found that the visual end was strongly activated, with a gradual transition to deactivation towards the auditory end of temporal G2. While the context task more strongly activated the visual end of G2 (than emotion), the emotion task activated bins towards the auditory end of this gradient (bins 3-4).

One finding that emerged consistently was the shift on G2 away from the auditory to the visual end across all four conditions probed. The two studies in this investigation were both visually presented, but with different modalities (verbal words versus pictures), yet produced a similar profile – especially with regard to the strong engagement of the visual end of G2. This finding is interesting, especially with regard to the prediction that verbal semantic tasks might recruit regions with connectivity to auditory more than visual processing streams, even when presented visually (Lambon Ralph et al., 2017; Visser et al., 2012). The differences across our two studies (i.e., verbal judgments versus generation to pictures) are also interesting. For example, a recent study found that feature matching leveraged visual spoke regions, while associative judgments showed stronger activation of the putative heteromodal hub (Chiou et al., 2018), yet we did not uncover a greater response for feature than associative judgments at the visual end of G2 (indeed, the opposite was true). The feature task did deactivate the auditory end of G2, with activation for this task focussed on the visual end of the gradient. It is notable that the associative content engaged the visual end of this gradient more strongly than the feature matching task (contrary to our prediction), but also that, despite differences in magnitude, *both* associative and feature judgments had strongest activation at the visual end of this gradient. Similarly, activation for both context and emotion generation was strongest at the visual end, although there were differences along the gradient, with emotion generation eliciting stronger responses in the middle-to-auditory end, while context generation showed stronger activation at the visual end.

While the profile of G2 was similar across both studies, the profile of G1 was not. Activation for feature and association judgments to verbal stimuli changed along the gradient in an orderly manner, however, this was not the case for emotion and context generation to pictures, which was characterised by deactivation at both extreme ends of G1, and activation across the middle to top (i.e., up to bin 9 of 10). This is interesting, because while picture semantic tasks are thought to constrain conceptual processing (e.g., they contain the visual features of the concepts, while verbal stimuli do not; Fernandino, Tong, Conant, Humphries, & Binder, 2022), the unimodal end of G1 deactivated and activation persisted across the middle to top of this gradient (i.e., into heteromodal cortex), despite the strong visual instantiation of the concepts, suggesting that participants engaged in some form of abstract semantic processing removed from the strong visual input. Furthermore, deactivation of the abstract emotion task in bins with strong connectivity to unimodal systems suggests that while emotion concepts can be grounded in sensorimotor systems (Barsalou, 2008; Niedenthal, 2007; Niedenthal et al., 2009; Vigliocco et al., 2014), they might also involve some abstraction away from these systems (Balgova et al., 2022; Mahon & Caramazza, 2008; Patterson, Nestor, & Rogers, 2007). However, while this finding aligns with a hub-like response for abstract conceptual processing, the decreased response at the top end of G1 for associative and feature judgments demonstrates the engagement of sensorimotor networks when making semantic judgments, decreasing gradually towards heteromodal cortex – a finding which is in line with both embodied accounts (increased unimodal response; Barsalou, 2008; Martin, 2016; Niedenthal, 2007) and graded accounts (the linear change in activation across the gradient; Bajada et al., 2017; Jackson et al., 2018; Lambon Ralph et al., 2017) of semantic cognition.

These complex findings highlight the need for future studies to use multiple presentation domains (e.g., auditory, visual etc), within the same participants, and across different process (e.g., judgments, internal generation, etc) and content (e.g., associative, feature, combinatorial, etc) types and modality (e.g., picture, verbal) to increase our understanding of the organisation of the temporal lobe, especially now that improved imaging techniques are available (e.g., MBME; Embleton et al., 2010; Halai et al., 2015; Halai et al., 2014; Poser & Norris, 2007; Poser et al., 2006). This investigation was limited in its ability to untangle unimodal responses due to stimulus presentation in only the visual domain, and was also unable to disentangle modality of the stimulus (e.g., words versus pictures) given the two studies differed in the task requirements (i.e., judgments versus generation). Furthermore, the modality associated with content of the task and/or concept was not varied in the verbal domain: we were unable to assess whether there was a stronger visual response for visual feature selection, compared to, e.g., auditory and/or motor feature selection, which might leverage sensorimotor codes. Ideally future studies would probe different semantic (and non-semantic) content across presentation domains – keeping the task constant to avoid confounds associated with task specific processes. However, it will also be important to start to untangle how task demands might influence engagement across the temporal lobe.

As Persichetti and colleagues (2021) noted, the candidate hub region for the graded hub model has shifted over time, as our understanding of the temporal lobe has progressed from coarse definitions based on cortical atrophy, to more recent imaging using protocols that maximise signal in this notoriously “tricky” region. While some progress has been made, based on meta-analyses and functional imaging studies (e.g., Balgova et al., 2022; Jackson et al., 2018; Rice, Hoffman, & Lambon Ralph, 2015), the debate over the location and even existence of a heteromodal hub continues (e.g., Lambon Ralph et al., 2017; Martin, 2016; Persichetti et al., 2021). This investigation did not set out to identify a candidate hub region (or indeed confirm or refute its existence), but rather examined how whole-brain connectivity gradients might contribute to our understanding of functional activation across temporal cortex for a range of content types and processes (associative and feature judgments and context and emotion generation). Given the between-subjects nature of this investigation, paired with the different task formats, this study is only a starting point from which to continue to probe our understanding of this complex region.

By leveraging different semantic content (associative, feature, emotion, context) and process (matching, generation), we investigated functional transitions along the temporal lobe, demonstrating a response that is at times graded when moving between unimodal and heteromodal temporal cortex (e.g., for associative and feature matching), and at others not (i.e., for internal generation to pictures: deactivation of extreme ends; non-graded activation across middle to heteromodal temporal cortex). The transitions from auditory to visual processing across temporal G2 showed a clear visual-bias regardless of content or process. These results highlight the complex nature of temporal lobe function and the need for more within-subjects’ investigations that probe multiple semantic domains and content to better understand how the temporal lobe is organised.

## Funding

The study was funded by the European Research Council [FLEXSEM-771863], and X. W. discloses support for the research of this work from National Natural Science Foundation of China (Grant No. 32300881) and Scientific Foundation of Institute of Psychology, Chinese Academy of Sciences (Grant Number. E1CX4725CX).

## Supplementary Materials

**Figure S1:**
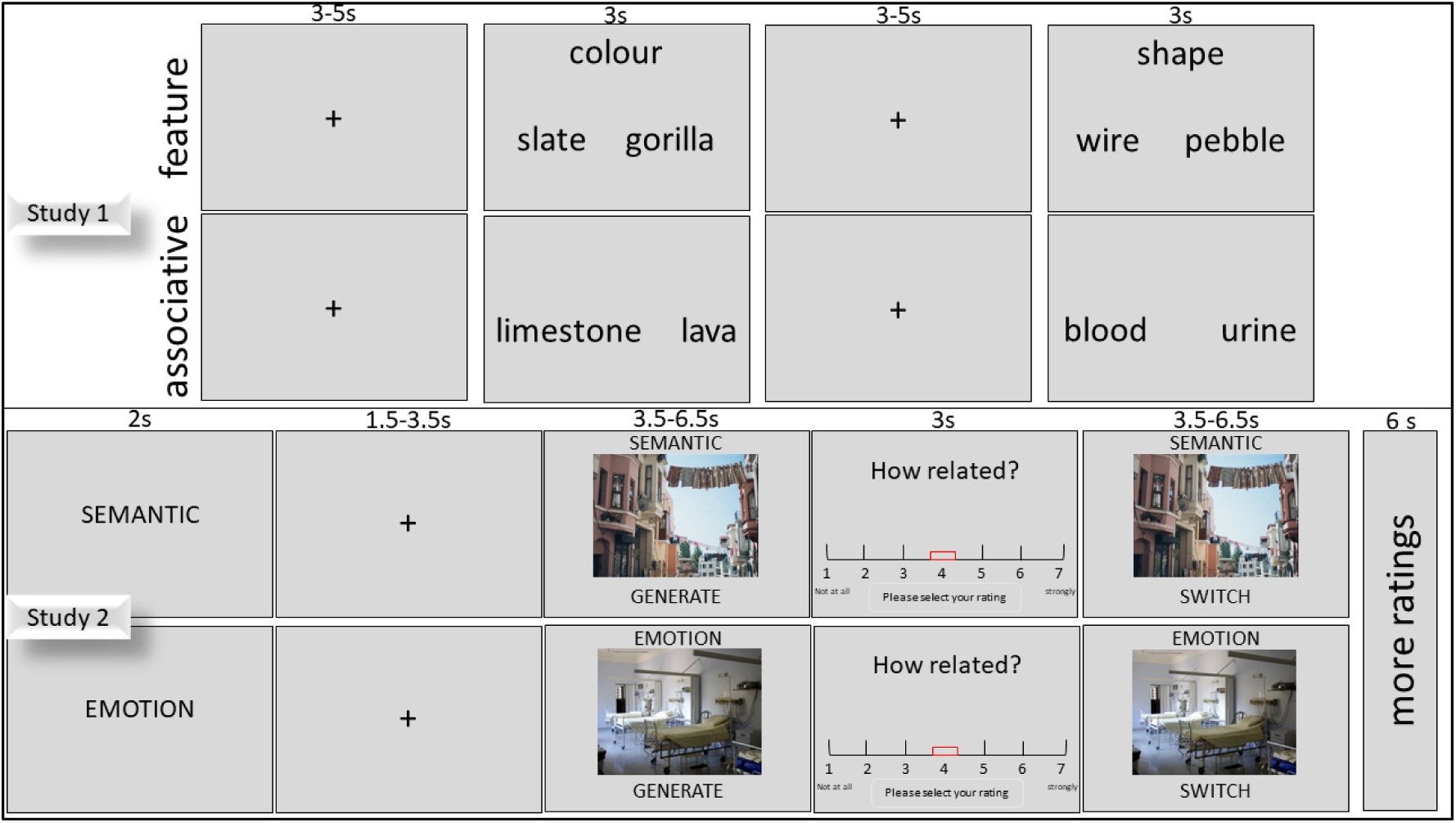
Task schematic for Study 1 (top) and Study 2 (bottom).

**Supplementary Table S1.**
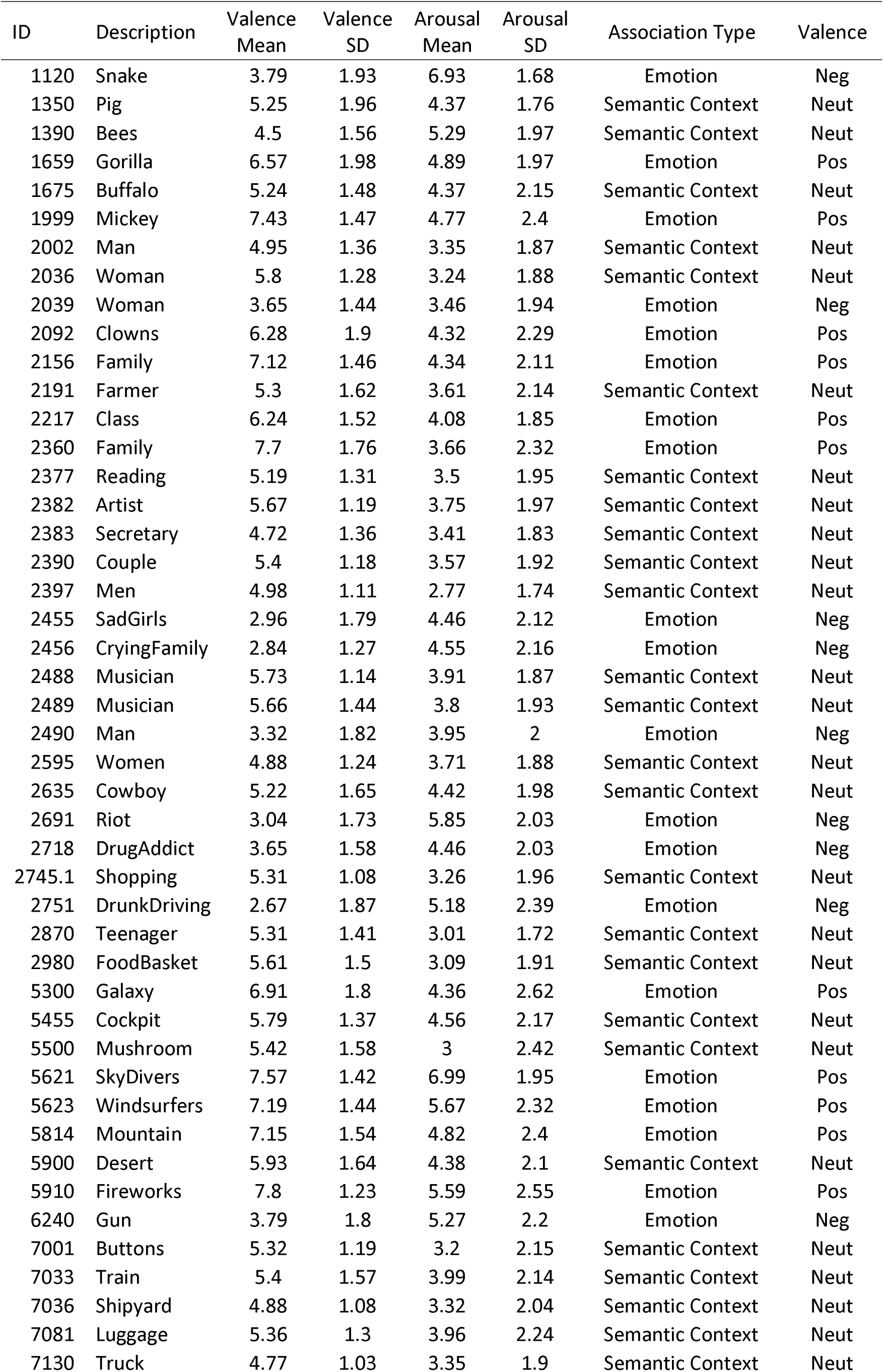
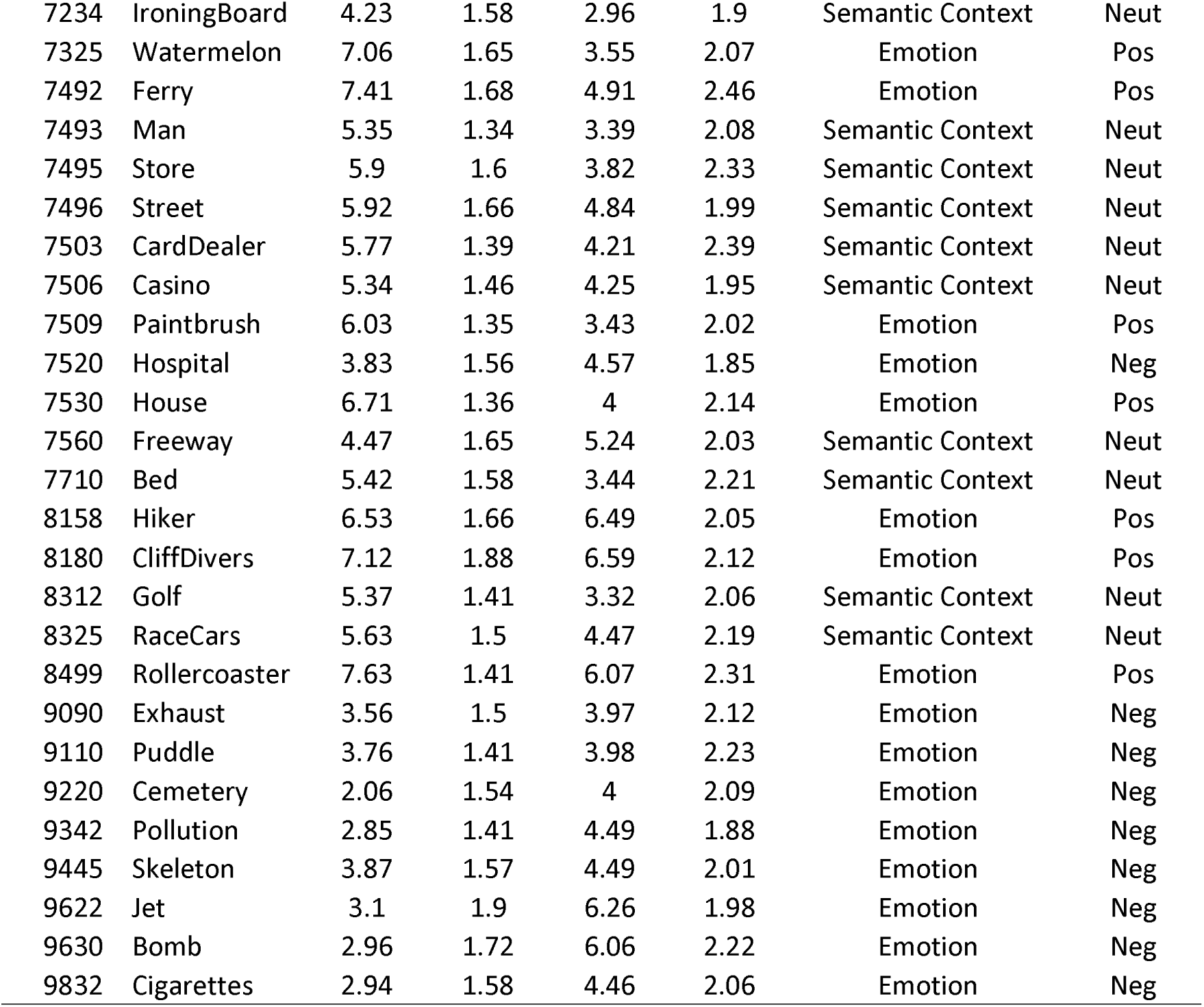
Identifiers for stimuli taken from the International Affective Picture System with mean and SD of valence and arousal ratings, and allocation of association type and categorical valence in the current study. Neg = Negative, Neut = Neutral, Pos = Positive.

**Table S2:**
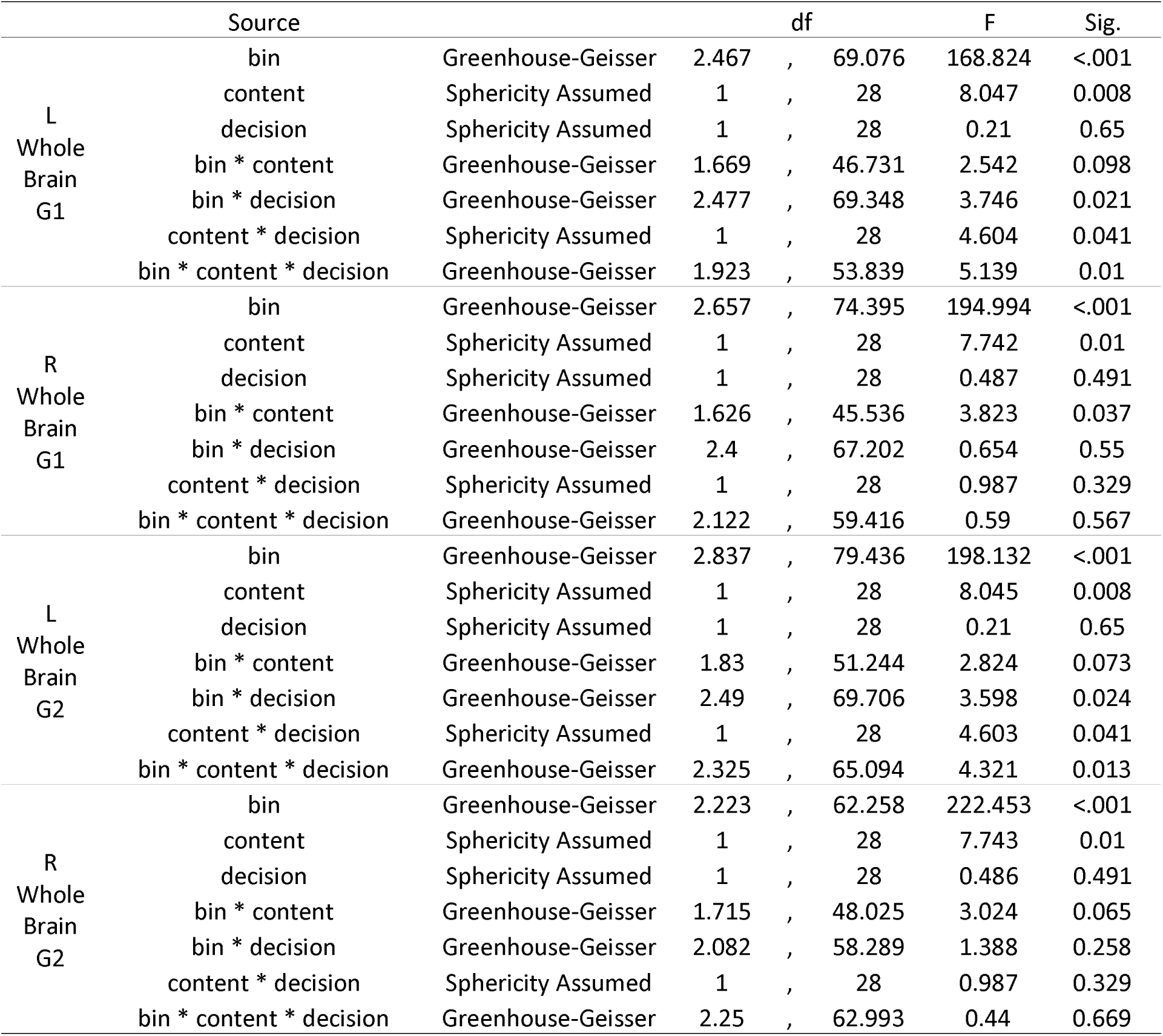
Whole Brain Gradients for Associative and Feature judgments.

**Table S3:**
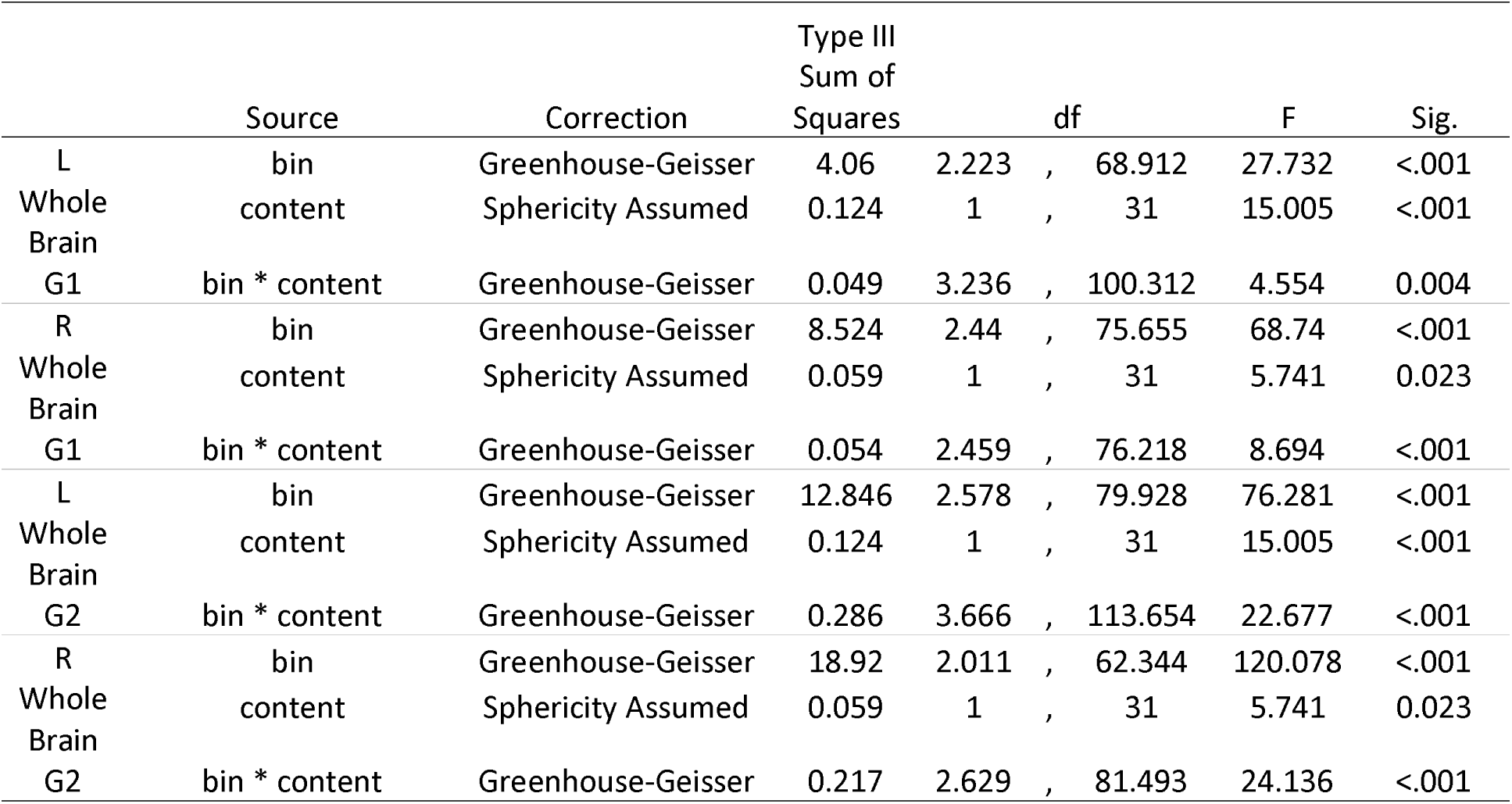
Whole Brain Gradients for Contex and Emotion Generation.

**Table S4:**
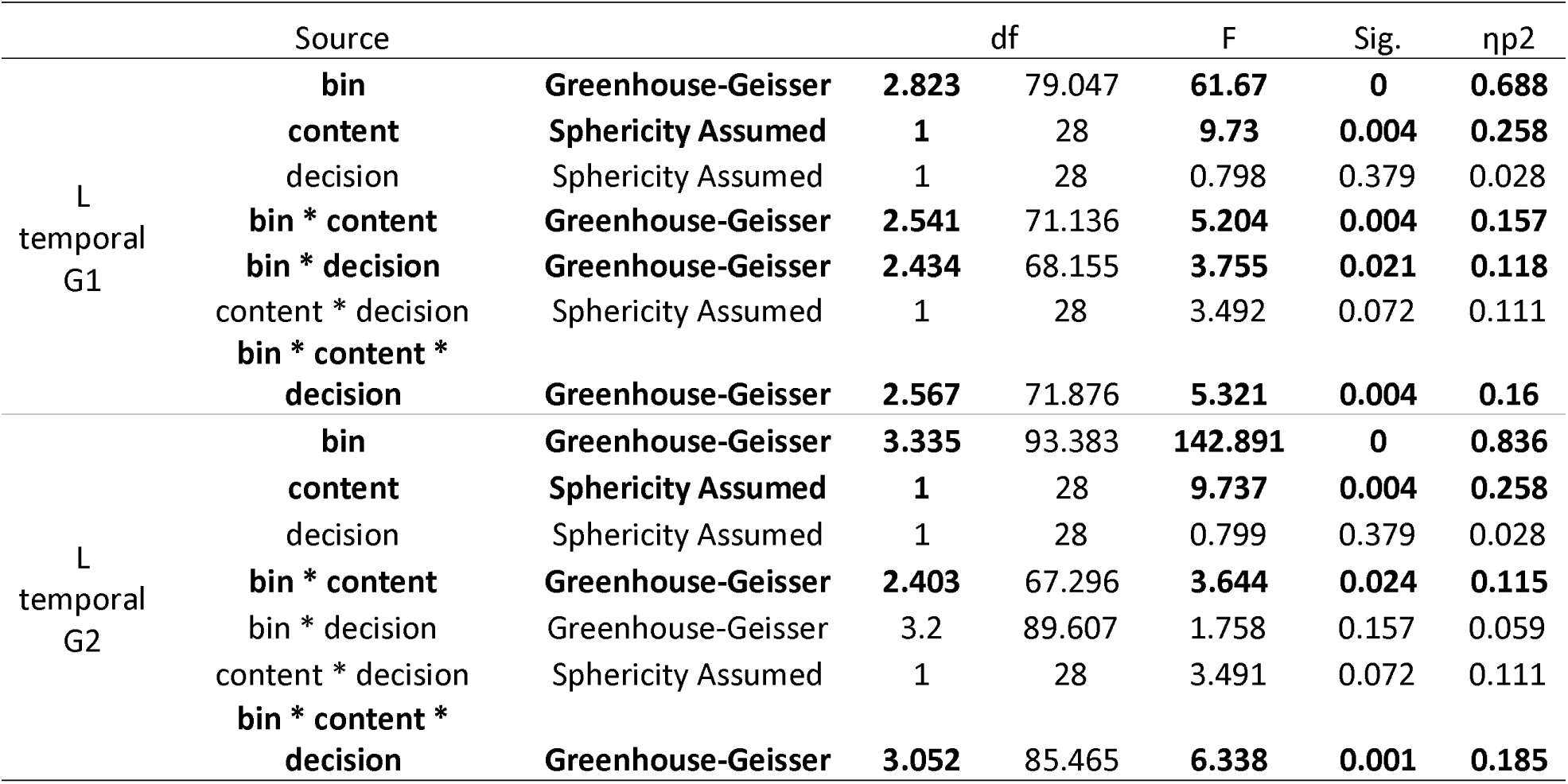
L temporal lobe G1 and G2 ANOVA’s.

**Table S5:**
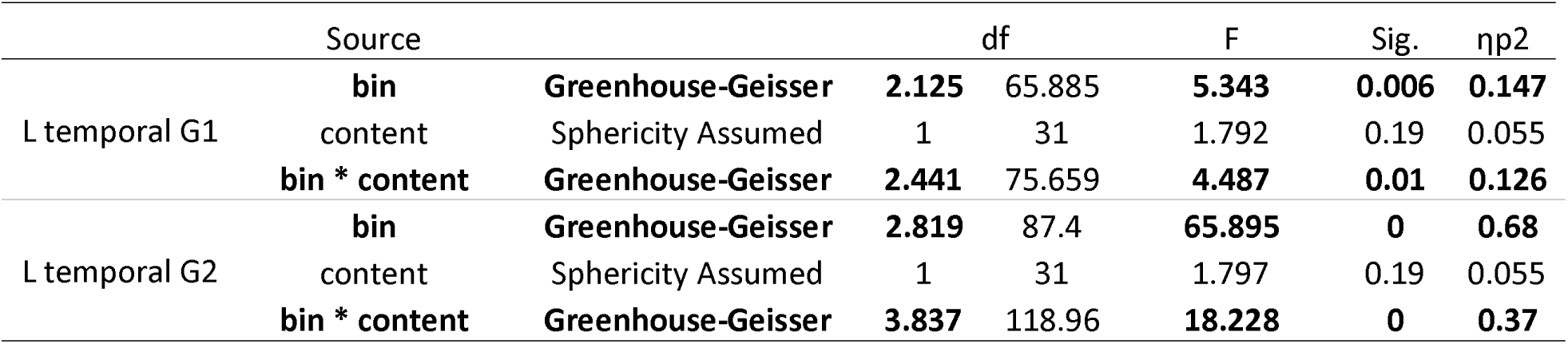
L temporal lobe G1 and G2 ANOVA’s.

**Table S6:**
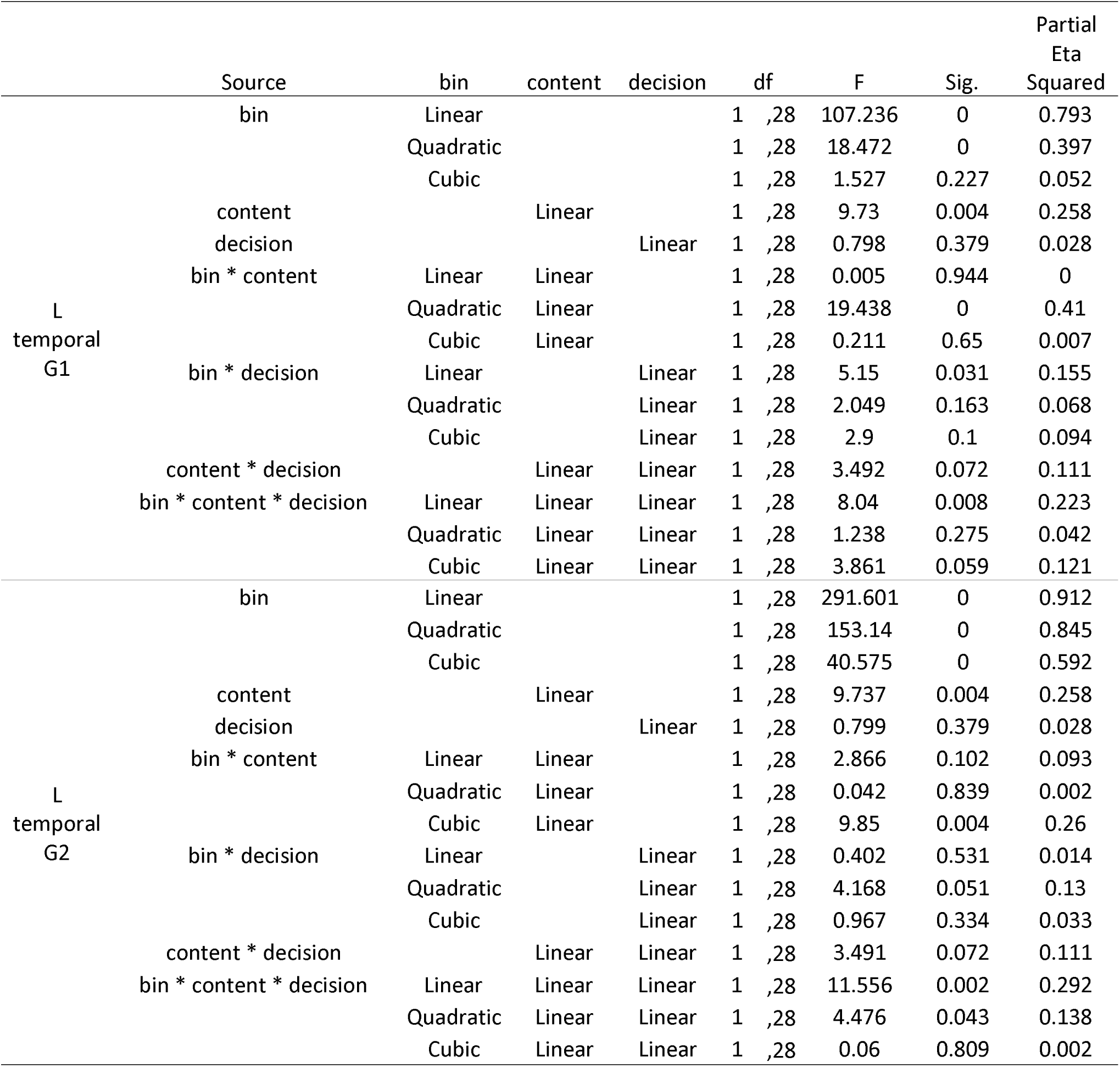
ANOVA trends for association and feature judgments.

**Table S7:**
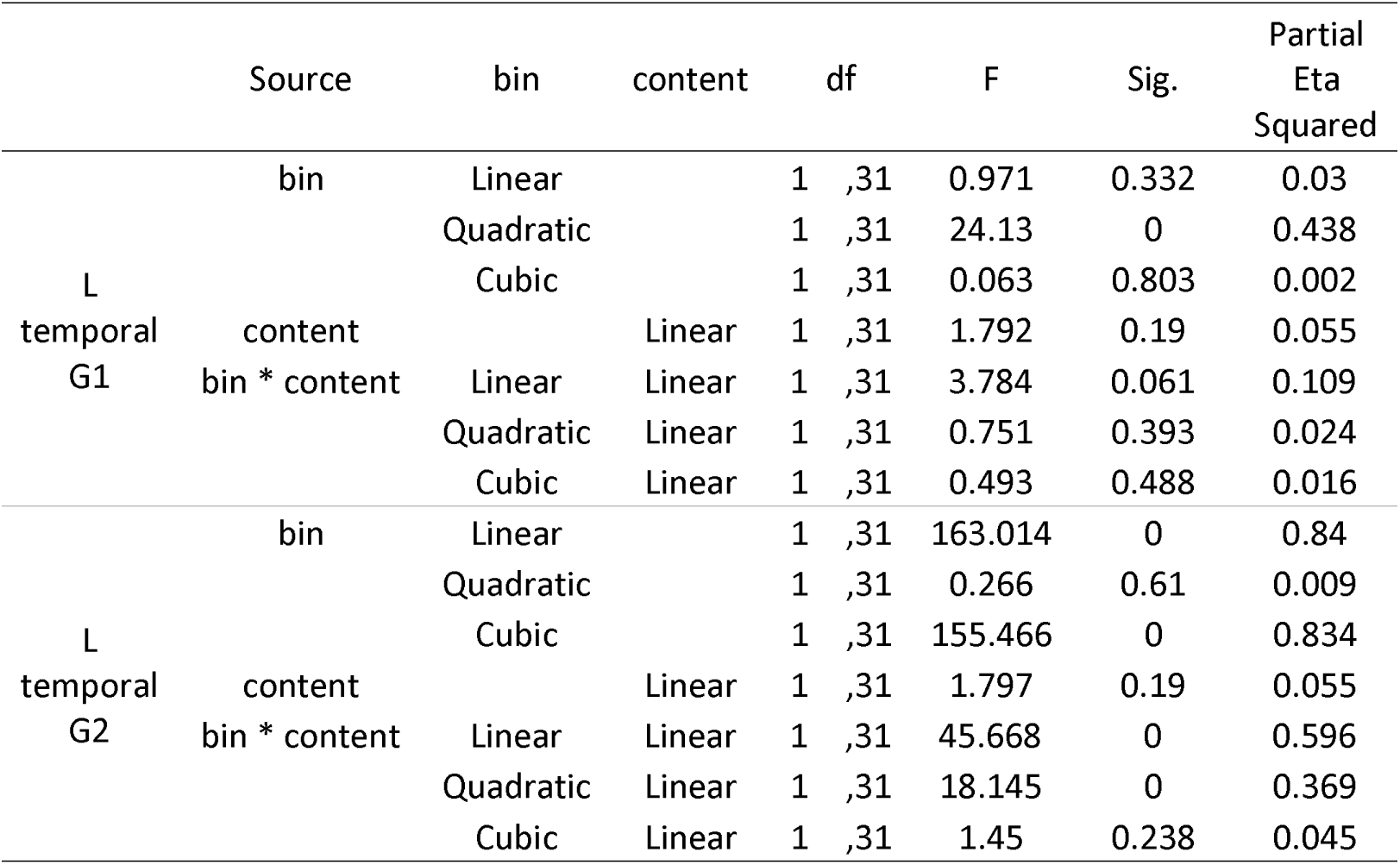
ANOVA trends for context and emotion generation.

**Table S8:**
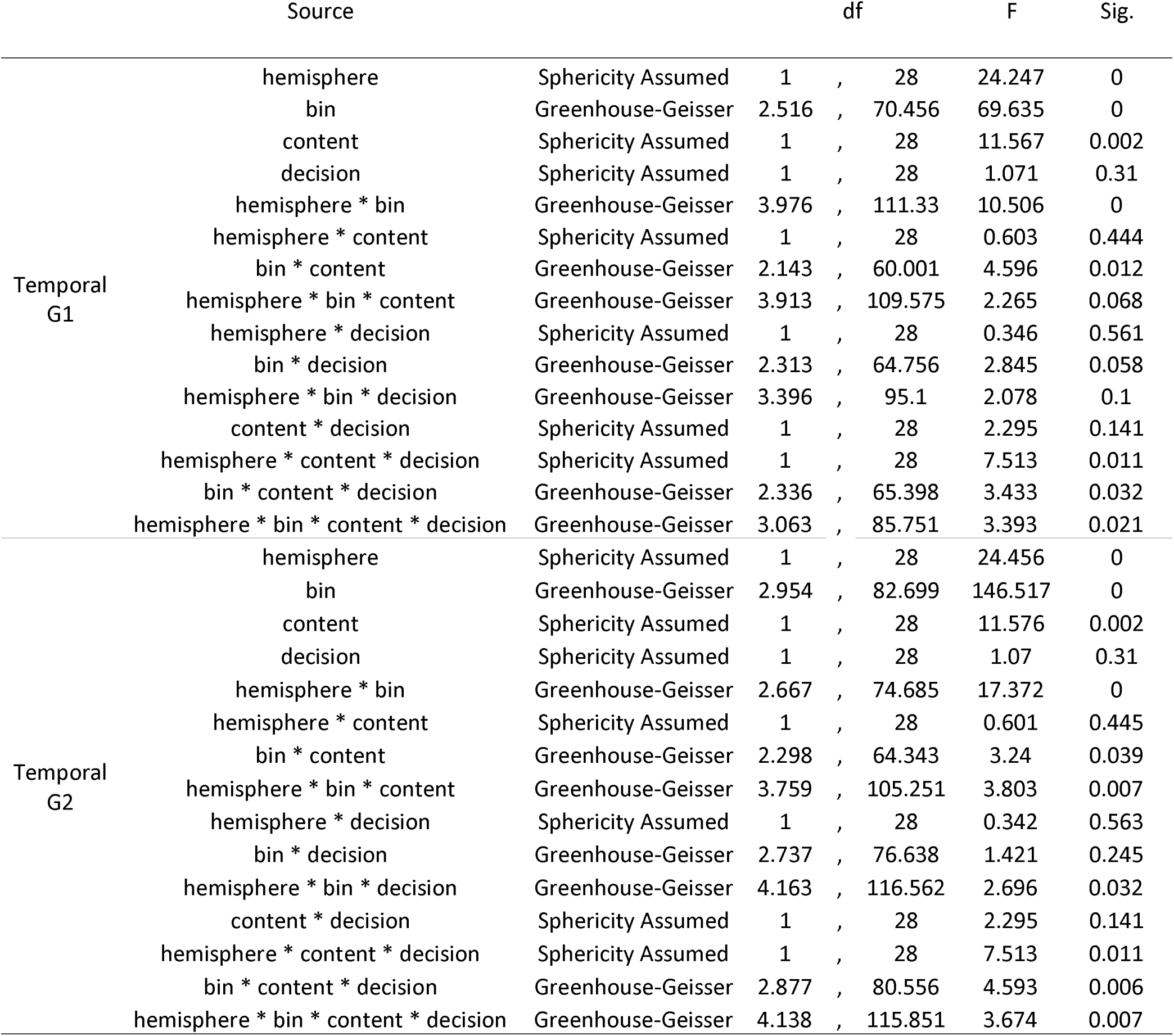
Left versus right temporal G1 and G2 for associative and feature judgments.

|  | Source |  | df |  | F | Sig. |
| --- | --- | --- | --- | --- | --- | --- |
| Temporal<br>G1 | hemisphere | Sphericity Assumed | 1 | , 28 | 24.247 | 0 |
|  | bin | Greenhouse-Geisser | 2.516 | , 70.456 | 69.635 | 0 |
|  | content | Sphericity Assumed | 1 | , 28 | 11.567 | 0.002 |
|  | decision | Sphericity Assumed | 1 | , 28 | 1.071 | 0.31 |
|  | hemisphere * bin | Greenhouse-Geisser | 3.976 | , 111.33 | 10.506 | 0 |
|  | hemisphere * content | Sphericity Assumed | 1 | , 28 | 0.603 | 0.444 |
|  | bin * content | Greenhouse-Geisser | 2.143 | , 60.001 | 4.596 | 0.012 |
|  | hemisphere * bin * content | Greenhouse-Geisser | 3.913 | , 109.575 | 2.265 | 0.068 |
|  | hemisphere * decision | Sphericity Assumed | 1 | , 28 | 0.346 | 0.561 |
|  | bin * decision | Greenhouse-Geisser | 2.313 | , 64.756 | 2.845 | 0.058 |
|  | hemisphere * bin * decision | Greenhouse-Geisser | 3.396 | , 95.1 | 2.078 | 0.1 |
|  | content * decision | Sphericity Assumed | 1 | , 28 | 2.295 | 0.141 |
|  | hemisphere * content * decision | Sphericity Assumed | 1 | , 28 | 7.513 | 0.011 |
|  | bin * content * decision | Greenhouse-Geisser | 2.336 | , 65.398 | 3.433 | 0.032 |
|  | hemisphere * bin * content * decision | Greenhouse-Geisser | 3.063 | , 85.751 | 3.393 | 0.021 |
| Temporal<br>G2 | hemisphere | Sphericity Assumed | 1 | , 28 | 24.456 | 0 |
|  | bin | Greenhouse-Geisser | 2.954 | , 82.699 | 146.517 | 0 |
|  | content | Sphericity Assumed | 1 | , 28 | 11.576 | 0.002 |
|  | decision | Sphericity Assumed | 1 | , 28 | 1.07 | 0.31 |
|  | hemisphere * bin | Greenhouse-Geisser | 2.667 | , 74.685 | 17.372 | 0 |
|  | hemisphere * content | Sphericity Assumed | 1 | , 28 | 0.601 | 0.445 |
|  | bin * content | Greenhouse-Geisser | 2.298 | , 64.343 | 3.24 | 0.039 |
|  | hemisphere * bin * content | Greenhouse-Geisser | 3.759 | , 105.251 | 3.803 | 0.007 |
|  | hemisphere * decision | Sphericity Assumed | 1 | , 28 | 0.342 | 0.563 |
|  | bin * decision | Greenhouse-Geisser | 2.737 | , 76.638 | 1.421 | 0.245 |
|  | hemisphere * bin * decision | Greenhouse-Geisser | 4.163 | , 116.562 | 2.696 | 0.032 |
|  | content * decision | Sphericity Assumed | 1 | , 28 | 2.295 | 0.141 |
|  | hemisphere * content * decision | Sphericity Assumed | 1 | , 28 | 7.513 | 0.011 |
|  | bin * content * decision | Greenhouse-Geisser | 2.877 | , 80.556 | 4.593 | 0.006 |
|  | hemisphere * bin * content * decision | Greenhouse-Geisser | 4.138 | , 115.851 | 3.674 | 0.007 |

**Table S9:**
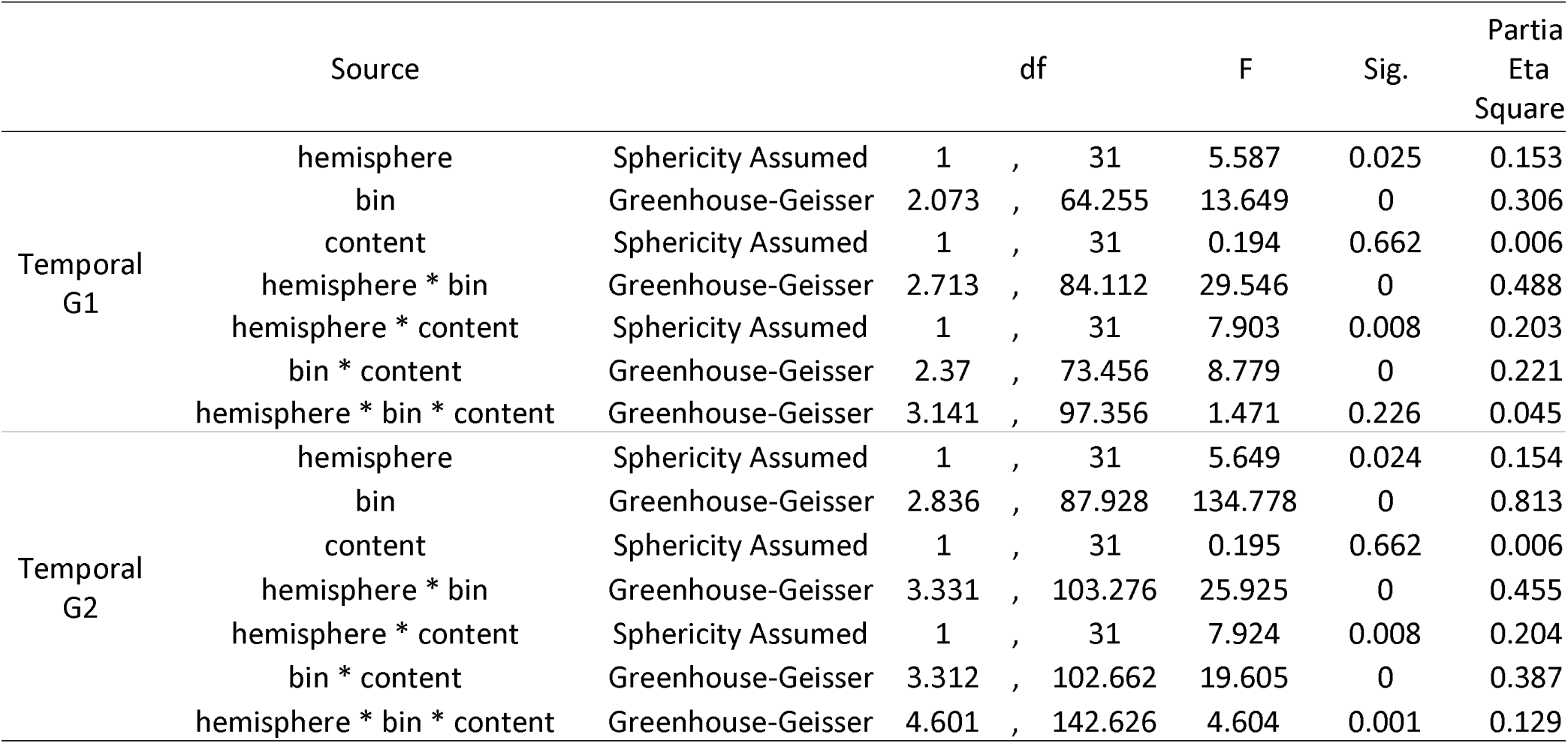
Left versus right temporal G1 and G2 for context and emotion generation.

**Table S10:**
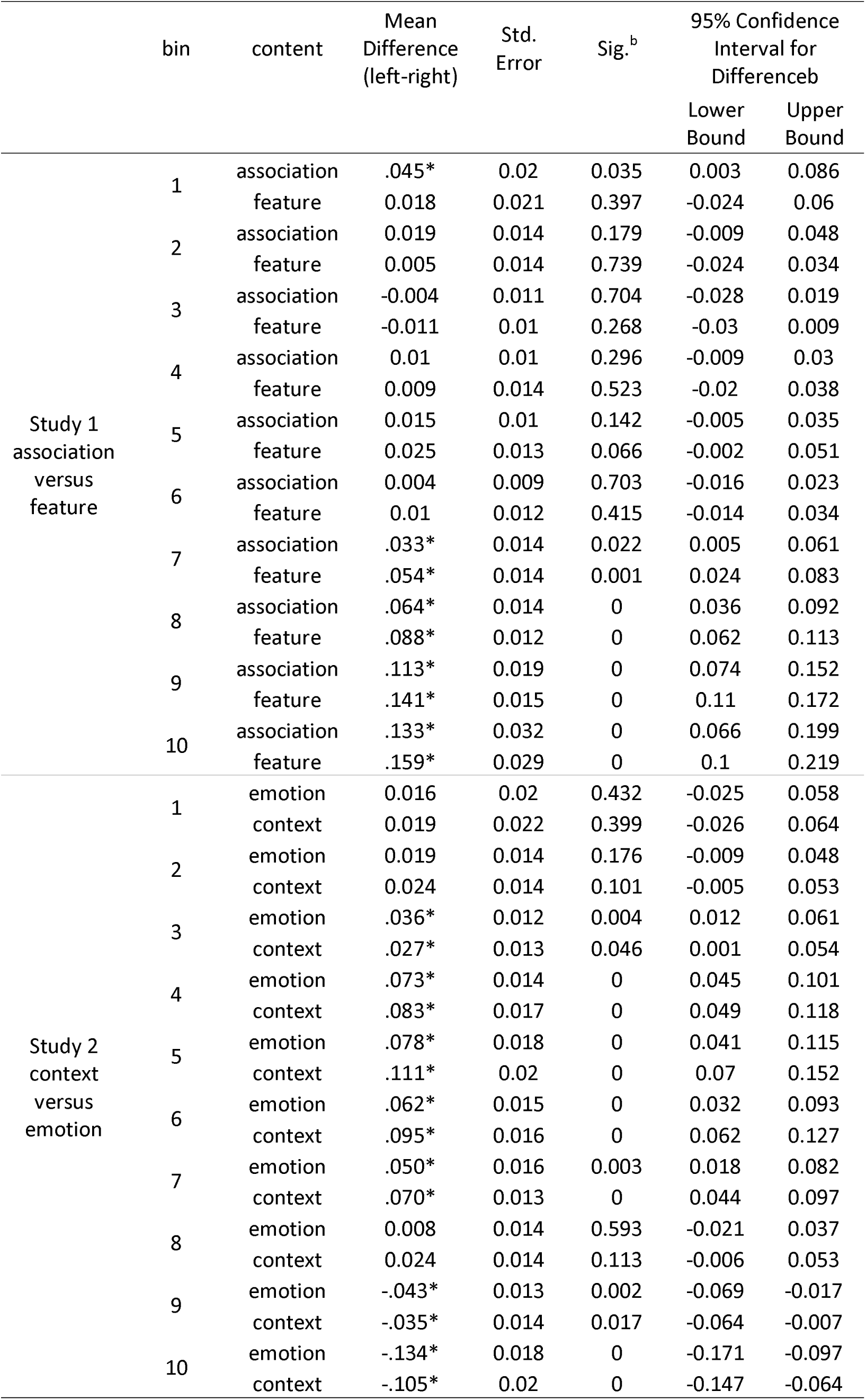
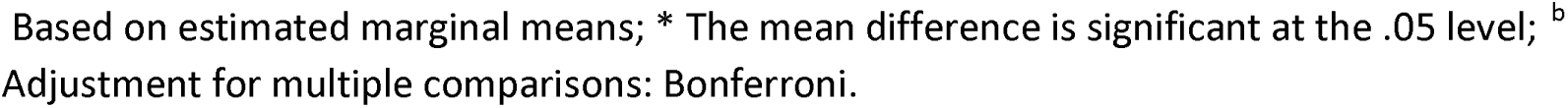
Left versus right temporal lobe G2 Bonferroni post-hoc tests.

## Notes

### Competing Interest Statement

The authors have declared no competing interest.

### Summary of Updates

This version of the manuscript has ben revised to update the introduction, methods, results and discussion following reviewer comments.

