## SupplementaryInformation for "Graded and sharp transitions in semantic function in left temporal lobe"

| ID | Description | Valence Mean | | Valence SD | Arousal Mean | Arousal SD | Association Type | Valence |
| --- | --- | --- | --- | --- | --- | --- | --- | --- |
| 1120 | Snake | | 3.79 | 1.93 | 6.93 | 1.68 | Emotion | Neg |
| 1350 | Pig | | 5.25 | 1.96 | 4.37 | 1.76 | Semantic Context | Neut |
| 1390 | Bees | | 4.5 | 1.56 | 5.29 | 1.97 | Semantic Context | Neut |
| 1659 | Gorilla | | 6.57 | 1.98 | 4.89 | 1.97 | Emotion | Pos |
| 1675 | Buffalo | | 5.24 | 1.48 | 4.37 | 2.15 | Semantic Context | Neut |
| 1999 | Mickey | | 7.43 | 1.47 | 4.77 | 2.4 | Emotion | Pos |
| 2002 | Man | | 4.95 | 1.36 | 3.35 | 1.87 | Semantic Context | Neut |
| 2036 | Woman | | 5.8 | 1.28 | 3.24 | 1.88 | Semantic Context | Neut |
| 2039 | Woman | | 3.65 | 1.44 | 3.46 | 1.94 | Emotion | Neg |
| 2092 | Clowns | | 6.28 | 1.9 | 4.32 | 2.29 | Emotion | Pos |
| 2156 | Family | | 7.12 | 1.46 | 4.34 | 2.11 | Emotion | Pos |
| 2191 | Farmer | | 5.3 | 1.62 | 3.61 | 2.14 | Semantic Context | Neut |
| 2217 | Class | | 6.24 | 1.52 | 4.08 | 1.85 | Emotion | Pos |
| 2360 | Family | | 7.7 | 1.76 | 3.66 | 2.32 | Emotion | Pos |
| 2377 | Reading | | 5.19 | 1.31 | 3.5 | 1.95 | Semantic Context | Neut |
| 2382 | Artist | | 5.67 | 1.19 | 3.75 | 1.97 | Semantic Context | Neut |
| 2383 | Secretary | | 4.72 | 1.36 | 3.41 | 1.83 | Semantic Context | Neut |
| 2390 | Couple | | 5.4 | 1.18 | 3.57 | 1.92 | Semantic Context | Neut |
| 2397 | Men | | 4.98 | 1.11 | 2.77 | 1.74 | Semantic Context | Neut |
| 2455 | SadGirls | | 2.96 | 1.79 | 4.46 | 2.12 | Emotion | Neg |
| 2456 | CryingFamily | | 2.84 | 1.27 | 4.55 | 2.16 | Emotion | Neg |
| 2488 | Musician | | 5.73 | 1.14 | 3.91 | 1.87 | Semantic Context | Neut |
| 2489 | Musician | | 5.66 | 1.44 | 3.8 | 1.93 | Semantic Context | Neut |
| 2490 | Man | | 3.32 | 1.82 | 3.95 | 2 | Emotion | Neg |
| 2595 | Women | | 4.88 | 1.24 | 3.71 | 1.88 | Semantic Context | Neut |
| 2635 | Cowboy | | 5.22 | 1.65 | 4.42 | 1.98 | Semantic Context | Neut |
| 2691 | Riot | | 3.04 | 1.73 | 5.85 | 2.03 | Emotion | Neg |
| 2718 | DrugAddict | | 3.65 | 1.58 | 4.46 | 2.03 | Emotion | Neg |
| 2745.1 | Shopping | | 5.31 | 1.08 | 3.26 | 1.96 | Semantic Context | Neut |
| 2751 | DrunkDriving | | 2.67 | 1.87 | 5.18 | 2.39 | Emotion | Neg |
| 2870 | Teenager | | 5.31 | 1.41 | 3.01 | 1.72 | Semantic Context | Neut |
| 2980 | FoodBasket | | 5.61 | 1.5 | 3.09 | 1.91 | Semantic Context | Neut |
| 5300 | Galaxy | | 6.91 | 1.8 | 4.36 | 2.62 | Emotion | Pos |
| 5455 | Cockpit | | 5.79 | 1.37 | 4.56 | 2.17 | Semantic Context | Neut |
| 5500 | Mushroom | | 5.42 | 1.58 | 3 | 2.42 | Semantic Context | Neut |
| 5621 | SkyDivers | | 7.57 | 1.42 | 6.99 | 1.95 | Emotion | Pos |
| 5623 | Windsurfers | | 7.19 | 1.44 | 5.67 | 2.32 | Emotion | Pos |
| 5814 | Mountain | | 7.15 | 1.54 | 4.82 | 2.4 | Emotion | Pos |
| 5900 | Desert | | 5.93 | 1.64 | 4.38 | 2.1 | Semantic Context | Neut |
| 5910 | Fireworks | | 7.8 | 1.23 | 5.59 | 2.55 | Emotion | Pos |
| 6240 | Gun | | 3.79 | 1.8 | 5.27 | 2.2 | Emotion | Neg |
| 7001 | Buttons | | 5.32 | 1.19 | 3.2 | 2.15 | Semantic Context | Neut |
| 7033 | Train | | 5.4 | 1.57 | 3.99 | 2.14 | Semantic Context | Neut |
| 7036 | Shipyard | | 4.88 | 1.08 | 3.32 | 2.04 | Semantic Context | Neut |
| 7081 | Luggage | | 5.36 | 1.3 | 3.96 | 2.24 | Semantic Context | Neut |
| 7130 | Truck | | 4.77 | 1.03 | 3.35 | 1.9 | Semantic Context | Neut |
| 7234 | IroningBoard | | 4.23 | 1.58 | 2.96 | 1.9 | Semantic Context | Neut |
| 7325 | Watermelon | | 7.06 | 1.65 | 3.55 | 2.07 | Emotion | Pos |
| 7492 | Ferry | | 7.41 | 1.68 | 4.91 | 2.46 | Emotion | Pos |
| 7493 | Man | | 5.35 | 1.34 | 3.39 | 2.08 | Semantic Context | Neut |
| 7495 | Store | | 5.9 | 1.6 | 3.82 | 2.33 | Semantic Context | Neut |
| 7496 | Street | | 5.92 | 1.66 | 4.84 | 1.99 | Semantic Context | Neut |
| 7503 | CardDealer | | 5.77 | 1.39 | 4.21 | 2.39 | Semantic Context | Neut |
| 7506 | Casino | | 5.34 | 1.46 | 4.25 | 1.95 | Semantic Context | Neut |
| 7509 | Paintbrush | | 6.03 | 1.35 | 3.43 | 2.02 | Emotion | Pos |
| 7520 | Hospital | | 3.83 | 1.56 | 4.57 | 1.85 | Emotion | Neg |
| 7530 | House | | 6.71 | 1.36 | 4 | 2.14 | Emotion | Pos |
| 7560 | Freeway | | 4.47 | 1.65 | 5.24 | 2.03 | Semantic Context | Neut |
| 7710 | Bed | | 5.42 | 1.58 | 3.44 | 2.21 | Semantic Context | Neut |
| 8158 | Hiker | | 6.53 | 1.66 | 6.49 | 2.05 | Emotion | Pos |
| 8180 | CliffDivers | | 7.12 | 1.88 | 6.59 | 2.12 | Emotion | Pos |
| 8312 | Golf | | 5.37 | 1.41 | 3.32 | 2.06 | Semantic Context | Neut |
| 8325 | RaceCars | | 5.63 | 1.5 | 4.47 | 2.19 | Semantic Context | Neut |
| 8499 | Rollercoaster | | 7.63 | 1.41 | 6.07 | 2.31 | Emotion | Pos |
| 9090 | Exhaust | | 3.56 | 1.5 | 3.97 | 2.12 | Emotion | Neg |
| 9110 | Puddle | | 3.76 | 1.41 | 3.98 | 2.23 | Emotion | Neg |
| 9220 | Cemetery | | 2.06 | 1.54 | 4 | 2.09 | Emotion | Neg |
| 9342 | Pollution | | 2.85 | 1.41 | 4.49 | 1.88 | Emotion | Neg |
| 9445 | Skeleton | | 3.87 | 1.57 | 4.49 | 2.01 | Emotion | Neg |
| 9622 | Jet | | 3.1 | 1.9 | 6.26 | 1.98 | Emotion | Neg |
| 9630 | Bomb | | 2.96 | 1.72 | 6.06 | 2.22 | Emotion | Neg |
| 9832 | Cigarettes | | 2.94 | 1.58 | 4.46 | 2.06 | Emotion | Neg |

Supplementary Table S1. Identifiers for stimuli taken from the International Affective Picture System with mean and SD of valence and arousal ratings, and allocation of association type and categorical valence in the current study. Neg = Negative, Neut = Neutral, Pos = Positive.

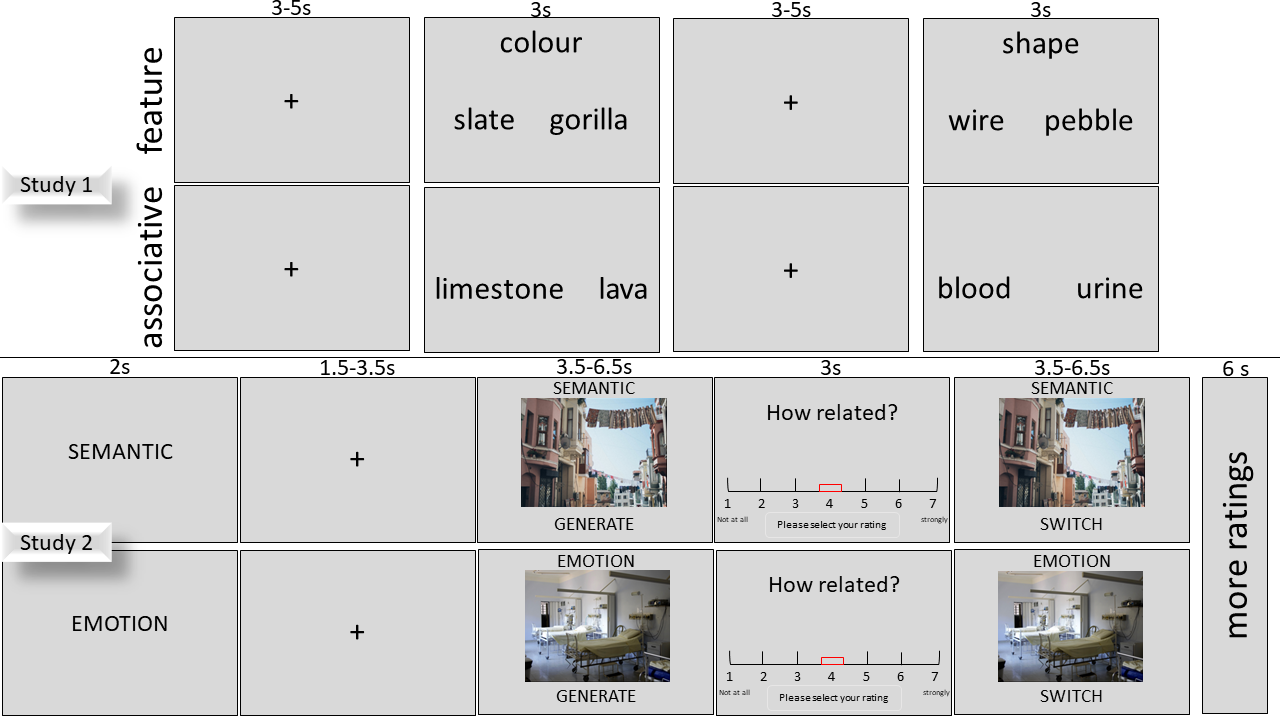

Figure S1: Task schematic for Study 1 (top) and Study 2 (bottom).

Table S2: Whole Brain Gradients for Associative and Feature judgments

|  | Source |  | df | | | F | Sig. |
| --- | --- | --- | --- | --- | --- | --- | --- |
| L Whole Brain G1 | bin | Greenhouse-Geisser | 2.467 | , | 69.076 | 168.824 | <.001 |
|  | content | Sphericity Assumed | 1 | , | 28 | 8.047 | 0.008 |
|  | decision | Sphericity Assumed | 1 | , | 28 | 0.21 | 0.65 |
|  | bin * content | Greenhouse-Geisser | 1.669 | , | 46.731 | 2.542 | 0.098 |
|  | bin * decision | Greenhouse-Geisser | 2.477 | , | 69.348 | 3.746 | 0.021 |
|  | content * decision | Sphericity Assumed | 1 | , | 28 | 4.604 | 0.041 |
|  | bin * content * decision | Greenhouse-Geisser | 1.923 | , | 53.839 | 5.139 | 0.01 |
| R Whole Brain G1 | bin | Greenhouse-Geisser | 2.657 | , | 74.395 | 194.994 | <.001 |
|  | content | Sphericity Assumed | 1 | , | 28 | 7.742 | 0.01 |
|  | decision | Sphericity Assumed | 1 | , | 28 | 0.487 | 0.491 |
|  | bin * content | Greenhouse-Geisser | 1.626 | , | 45.536 | 3.823 | 0.037 |
|  | bin * decision | Greenhouse-Geisser | 2.4 | , | 67.202 | 0.654 | 0.55 |
|  | content * decision | Sphericity Assumed | 1 | , | 28 | 0.987 | 0.329 |
|  | bin * content * decision | Greenhouse-Geisser | 2.122 | , | 59.416 | 0.59 | 0.567 |
| L Whole Brain G2 | bin | Greenhouse-Geisser | 2.837 | , | 79.436 | 198.132 | <.001 |
|  | content | Sphericity Assumed | 1 | , | 28 | 8.045 | 0.008 |
|  | decision | Sphericity Assumed | 1 | , | 28 | 0.21 | 0.65 |
|  | bin * content | Greenhouse-Geisser | 1.83 | , | 51.244 | 2.824 | 0.073 |
|  | bin * decision | Greenhouse-Geisser | 2.49 | , | 69.706 | 3.598 | 0.024 |
|  | content * decision | Sphericity Assumed | 1 | , | 28 | 4.603 | 0.041 |
|  | bin * content * decision | Greenhouse-Geisser | 2.325 | , | 65.094 | 4.321 | 0.013 |
| R Whole Brain G2 | bin | Greenhouse-Geisser | 2.223 | , | 62.258 | 222.453 | <.001 |
|  | content | Sphericity Assumed | 1 | , | 28 | 7.743 | 0.01 |
|  | decision | Sphericity Assumed | 1 | , | 28 | 0.486 | 0.491 |
|  | bin * content | Greenhouse-Geisser | 1.715 | , | 48.025 | 3.024 | 0.065 |
|  | bin * decision | Greenhouse-Geisser | 2.082 | , | 58.289 | 1.388 | 0.258 |
|  | content * decision | Sphericity Assumed | 1 | , | 28 | 0.987 | 0.329 |
|  | bin * content * decision | Greenhouse-Geisser | 2.25 | , | 62.993 | 0.44 | 0.669 |

Table S3: Whole Brain Gradients for Contex and Emotion Generation

|  | Source | Correction | Type III Sum of Squares | df | | | F | Sig. |
| --- | --- | --- | --- | --- | --- | --- | --- | --- |
| L Whole Brain G1 | bin | Greenhouse-Geisser | 4.06 | 2.223 | , | 68.912 | 27.732 | <.001 |
|  | content | Sphericity Assumed | 0.124 | 1 | , | 31 | 15.005 | <.001 |
|  | bin * content | Greenhouse-Geisser | 0.049 | 3.236 | , | 100.312 | 4.554 | 0.004 |
| R Whole Brain G1 | bin | Greenhouse-Geisser | 8.524 | 2.44 | , | 75.655 | 68.74 | <.001 |
|  | content | Sphericity Assumed | 0.059 | 1 | , | 31 | 5.741 | 0.023 |
|  | bin * content | Greenhouse-Geisser | 0.054 | 2.459 | , | 76.218 | 8.694 | <.001 |
| L Whole Brain G2 | bin | Greenhouse-Geisser | 12.846 | 2.578 | , | 79.928 | 76.281 | <.001 |
|  | content | Sphericity Assumed | 0.124 | 1 | , | 31 | 15.005 | <.001 |
|  | bin * content | Greenhouse-Geisser | 0.286 | 3.666 | , | 113.654 | 22.677 | <.001 |
| R Whole Brain G2 | bin | Greenhouse-Geisser | 18.92 | 2.011 | , | 62.344 | 120.078 | <.001 |
|  | content | Sphericity Assumed | 0.059 | 1 | , | 31 | 5.741 | 0.023 |
|  | bin * content | Greenhouse-Geisser | 0.217 | 2.629 | , | 81.493 | 24.136 | <.001 |

Table S4: L temporal lobe G1 and G2 ANOVA’s

|  | Source |  | df | | F | Sig. | ηp2 |
| --- | --- | --- | --- | --- | --- | --- | --- |
| L temporal G1 | **bin** | **Greenhouse-Geisser** | **2.823** | 79.047 | **61.67** | **0** | **0.688** |
|  | **content** | **Sphericity Assumed** | **1** | 28 | **9.73** | **0.004** | **0.258** |
|  | decision | Sphericity Assumed | 1 | 28 | 0.798 | 0.379 | 0.028 |
|  | **bin * content** | **Greenhouse-Geisser** | **2.541** | 71.136 | **5.204** | **0.004** | **0.157** |
|  | **bin * decision** | **Greenhouse-Geisser** | **2.434** | 68.155 | **3.755** | **0.021** | **0.118** |
|  | content * decision | Sphericity Assumed | 1 | 28 | 3.492 | 0.072 | 0.111 |
|  | **bin * content * decision** | **Greenhouse-Geisser** | **2.567** | 71.876 | **5.321** | **0.004** | **0.16** |
| L temporal G2 | **bin** | **Greenhouse-Geisser** | **3.335** | 93.383 | **142.891** | **0** | **0.836** |
|  | **content** | **Sphericity Assumed** | **1** | 28 | **9.737** | **0.004** | **0.258** |
|  | decision | Sphericity Assumed | 1 | 28 | 0.799 | 0.379 | 0.028 |
|  | **bin * content** | **Greenhouse-Geisser** | **2.403** | 67.296 | **3.644** | **0.024** | **0.115** |
|  | bin * decision | Greenhouse-Geisser | 3.2 | 89.607 | 1.758 | 0.157 | 0.059 |
|  | content * decision | Sphericity Assumed | 1 | 28 | 3.491 | 0.072 | 0.111 |
|  | **bin * content * decision** | **Greenhouse-Geisser** | **3.052** | 85.465 | **6.338** | **0.001** | **0.185** |

Table S5: L temporal lobe G1 and G2 ANOVA’s

|  | Source |  | df | | F | Sig. | ηp2 |
| --- | --- | --- | --- | --- | --- | --- | --- |
| L temporal G1 | **bin** | **Greenhouse-Geisser** | **2.125** | 65.885 | **5.343** | **0.006** | **0.147** |
|  | content | Sphericity Assumed | 1 | 31 | 1.792 | 0.19 | 0.055 |
|  | **bin * content** | **Greenhouse-Geisser** | **2.441** | 75.659 | **4.487** | **0.01** | **0.126** |
| L temporal G2 | **bin** | **Greenhouse-Geisser** | **2.819** | 87.4 | **65.895** | **0** | **0.68** |
|  | content | Sphericity Assumed | 1 | 31 | 1.797 | 0.19 | 0.055 |
|  | **bin * content** | **Greenhouse-Geisser** | **3.837** | 118.96 | **18.228** | **0** | **0.37** |

Table S6: ANOVA trends for association and feature judgments

|  | Source | bin | content | decision | df | | F | Sig. | Partial Eta Squared |
| --- | --- | --- | --- | --- | --- | --- | --- | --- | --- |
| L temporal G1 | bin | Linear |  |  | 1 | ,28 | 107.236 | 0 | 0.793 |
|  |  | Quadratic |  |  | 1 | ,28 | 18.472 | 0 | 0.397 |
|  |  | Cubic |  |  | 1 | ,28 | 1.527 | 0.227 | 0.052 |
|  | content |  | Linear |  | 1 | ,28 | 9.73 | 0.004 | 0.258 |
|  | decision |  |  | Linear | 1 | ,28 | 0.798 | 0.379 | 0.028 |
|  | bin * content | Linear | Linear |  | 1 | ,28 | 0.005 | 0.944 | 0 |
|  |  | Quadratic | Linear |  | 1 | ,28 | 19.438 | 0 | 0.41 |
|  |  | Cubic | Linear |  | 1 | ,28 | 0.211 | 0.65 | 0.007 |
|  | bin * decision | Linear |  | Linear | 1 | ,28 | 5.15 | 0.031 | 0.155 |
|  |  | Quadratic |  | Linear | 1 | ,28 | 2.049 | 0.163 | 0.068 |
|  |  | Cubic |  | Linear | 1 | ,28 | 2.9 | 0.1 | 0.094 |
|  | content * decision |  | Linear | Linear | 1 | ,28 | 3.492 | 0.072 | 0.111 |
|  | bin * content * decision | Linear | Linear | Linear | 1 | ,28 | 8.04 | 0.008 | 0.223 |
|  |  | Quadratic | Linear | Linear | 1 | ,28 | 1.238 | 0.275 | 0.042 |
|  |  | Cubic | Linear | Linear | 1 | ,28 | 3.861 | 0.059 | 0.121 |
| L temporal G2 | bin | Linear |  |  | 1 | ,28 | 291.601 | 0 | 0.912 |
|  |  | Quadratic |  |  | 1 | ,28 | 153.14 | 0 | 0.845 |
|  |  | Cubic |  |  | 1 | ,28 | 40.575 | 0 | 0.592 |
|  | content |  | Linear |  | 1 | ,28 | 9.737 | 0.004 | 0.258 |
|  | decision |  |  | Linear | 1 | ,28 | 0.799 | 0.379 | 0.028 |
|  | bin * content | Linear | Linear |  | 1 | ,28 | 2.866 | 0.102 | 0.093 |
|  |  | Quadratic | Linear |  | 1 | ,28 | 0.042 | 0.839 | 0.002 |
|  |  | Cubic | Linear |  | 1 | ,28 | 9.85 | 0.004 | 0.26 |
|  | bin * decision | Linear |  | Linear | 1 | ,28 | 0.402 | 0.531 | 0.014 |
|  |  | Quadratic |  | Linear | 1 | ,28 | 4.168 | 0.051 | 0.13 |
|  |  | Cubic |  | Linear | 1 | ,28 | 0.967 | 0.334 | 0.033 |
|  | content * decision |  | Linear | Linear | 1 | ,28 | 3.491 | 0.072 | 0.111 |
|  | bin * content * decision | Linear | Linear | Linear | 1 | ,28 | 11.556 | 0.002 | 0.292 |
|  |  | Quadratic | Linear | Linear | 1 | ,28 | 4.476 | 0.043 | 0.138 |
|  |  | Cubic | Linear | Linear | 1 | ,28 | 0.06 | 0.809 | 0.002 |

Table S7: ANOVA trends for context and emotion generation

|  | Source | bin | content | df | | F | Sig. | Partial Eta Squared |
| --- | --- | --- | --- | --- | --- | --- | --- | --- |
| L temporal G1 | bin | Linear |  | 1 | ,31 | 0.971 | 0.332 | 0.03 |
|  |  | Quadratic |  | 1 | ,31 | 24.13 | 0 | 0.438 |
|  |  | Cubic |  | 1 | ,31 | 0.063 | 0.803 | 0.002 |
|  | content |  | Linear | 1 | ,31 | 1.792 | 0.19 | 0.055 |
|  | bin * content | Linear | Linear | 1 | ,31 | 3.784 | 0.061 | 0.109 |
|  |  | Quadratic | Linear | 1 | ,31 | 0.751 | 0.393 | 0.024 |
|  |  | Cubic | Linear | 1 | ,31 | 0.493 | 0.488 | 0.016 |
| L temporal G2 | bin | Linear |  | 1 | ,31 | 163.014 | 0 | 0.84 |
|  |  | Quadratic |  | 1 | ,31 | 0.266 | 0.61 | 0.009 |
|  |  | Cubic |  | 1 | ,31 | 155.466 | 0 | 0.834 |
|  | content |  | Linear | 1 | ,31 | 1.797 | 0.19 | 0.055 |
|  | bin * content | Linear | Linear | 1 | ,31 | 45.668 | 0 | 0.596 |
|  |  | Quadratic | Linear | 1 | ,31 | 18.145 | 0 | 0.369 |
|  |  | Cubic | Linear | 1 | ,31 | 1.45 | 0.238 | 0.045 |

Table S8: Left versus right temporal G1 and G2 for associative and feature judgments

|  | Source |  | df | | | F | Sig. | Partial Eta Squared |
| --- | --- | --- | --- | --- | --- | --- | --- | --- |
| Temporal G1 | hemisphere | Sphericity Assumed | 1 | , | 28 | 24.247 | 0 | 0.464 |
|  | bin | Greenhouse-Geisser | 2.516 | , | 70.456 | 69.635 | 0 | 0.713 |
|  | content | Sphericity Assumed | 1 | , | 28 | 11.567 | 0.002 | 0.292 |
|  | decision | Sphericity Assumed | 1 | , | 28 | 1.071 | 0.31 | 0.037 |
|  | hemisphere * bin | Greenhouse-Geisser | 3.976 | , | 111.33 | 10.506 | 0 | 0.273 |
|  | hemisphere * content | Sphericity Assumed | 1 | , | 28 | 0.603 | 0.444 | 0.021 |
|  | bin * content | Greenhouse-Geisser | 2.143 | , | 60.001 | 4.596 | 0.012 | 0.141 |
|  | hemisphere * bin * content | Greenhouse-Geisser | 3.913 | , | 109.575 | 2.265 | 0.068 | 0.075 |
|  | hemisphere * decision | Sphericity Assumed | 1 | , | 28 | 0.346 | 0.561 | 0.012 |
|  | bin * decision | Greenhouse-Geisser | 2.313 | , | 64.756 | 2.845 | 0.058 | 0.092 |
|  | hemisphere * bin * decision | Greenhouse-Geisser | 3.396 | , | 95.1 | 2.078 | 0.1 | 0.069 |
|  | content * decision | Sphericity Assumed | 1 | , | 28 | 2.295 | 0.141 | 0.076 |
|  | hemisphere * content * decision | Sphericity Assumed | 1 | , | 28 | 7.513 | 0.011 | 0.212 |
|  | bin * content * decision | Greenhouse-Geisser | 2.336 | , | 65.398 | 3.433 | 0.032 | 0.109 |
|  | hemisphere * bin * content * decision | Greenhouse-Geisser | 3.063 | , | 85.751 | 3.393 | 0.021 | 0.108 |
| Temporal G2 | hemisphere | Sphericity Assumed | 1 | , | 28 | 24.456 | 0 | 0.466 |
|  | bin | Greenhouse-Geisser | 2.954 | , | 82.699 | 146.517 | 0 | 0.84 |
|  | content | Sphericity Assumed | 1 | , | 28 | 11.576 | 0.002 | 0.292 |
|  | decision | Sphericity Assumed | 1 | , | 28 | 1.07 | 0.31 | 0.037 |
|  | hemisphere * bin | Greenhouse-Geisser | 2.667 | , | 74.685 | 17.372 | 0 | 0.383 |
|  | hemisphere * content | Sphericity Assumed | 1 | , | 28 | 0.601 | 0.445 | 0.021 |
|  | bin * content | Greenhouse-Geisser | 2.298 | , | 64.343 | 3.24 | 0.039 | 0.104 |
|  | hemisphere * bin * content | Greenhouse-Geisser | 3.759 | , | 105.251 | 3.803 | 0.007 | 0.12 |
|  | hemisphere * decision | Sphericity Assumed | 1 | , | 28 | 0.342 | 0.563 | 0.012 |
|  | bin * decision | Greenhouse-Geisser | 2.737 | , | 76.638 | 1.421 | 0.245 | 0.048 |
|  | hemisphere * bin * decision | Greenhouse-Geisser | 4.163 | , | 116.562 | 2.696 | 0.032 | 0.088 |
|  | content * decision | Sphericity Assumed | 1 | , | 28 | 2.295 | 0.141 | 0.076 |
|  | hemisphere * content * decision | Sphericity Assumed | 1 | , | 28 | 7.513 | 0.011 | 0.212 |
|  | bin * content * decision | Greenhouse-Geisser | 2.877 | , | 80.556 | 4.593 | 0.006 | 0.141 |
|  | hemisphere * bin * content * decision | Greenhouse-Geisser | 4.138 | , | 115.851 | 3.674 | 0.007 | 0.116 |

Table S9: Left versus right temporal G1 and G2 for context and emotion generation

|  | Source |  | df | | | F | Sig. | Partial Eta Squared |
| --- | --- | --- | --- | --- | --- | --- | --- | --- |
| Temporal G1 | hemisphere | Sphericity Assumed | 1 | , | 31 | 5.587 | 0.025 | 0.153 |
|  | bin | Greenhouse-Geisser | 2.073 | , | 64.255 | 13.649 | 0 | 0.306 |
|  | content | Sphericity Assumed | 1 | , | 31 | 0.194 | 0.662 | 0.006 |
|  | hemisphere * bin | Greenhouse-Geisser | 2.713 | , | 84.112 | 29.546 | 0 | 0.488 |
|  | hemisphere * content | Sphericity Assumed | 1 | , | 31 | 7.903 | 0.008 | 0.203 |
|  | bin * content | Greenhouse-Geisser | 2.37 | , | 73.456 | 8.779 | 0 | 0.221 |
|  | hemisphere * bin * content | Greenhouse-Geisser | 3.141 | , | 97.356 | 1.471 | 0.226 | 0.045 |
| Temporal G2 | hemisphere | Sphericity Assumed | 1 | , | 31 | 5.649 | 0.024 | 0.154 |
|  | bin | Greenhouse-Geisser | 2.836 | , | 87.928 | 134.778 | 0 | 0.813 |
|  | content | Sphericity Assumed | 1 | , | 31 | 0.195 | 0.662 | 0.006 |
|  | hemisphere * bin | Greenhouse-Geisser | 3.331 | , | 103.276 | 25.925 | 0 | 0.455 |
|  | hemisphere * content | Sphericity Assumed | 1 | , | 31 | 7.924 | 0.008 | 0.204 |
|  | bin * content | Greenhouse-Geisser | 3.312 | , | 102.662 | 19.605 | 0 | 0.387 |
|  | hemisphere * bin * content | Greenhouse-Geisser | 4.601 | , | 142.626 | 4.604 | 0.001 | 0.129 |

Table S10: Left versus right temporal lobe G2 Bonferroni post-hoc tests

|  | bin | content | Mean Difference (left-right) | Std. Error | Sig.^b^ | 95% Confidence Interval for Differenceb | |
| --- | --- | --- | --- | --- | --- | --- | --- |
|  |  |  |  |  |  | Lower Bound | Upper Bound |
| Study 1 association versus feature | 1 | association | .045* | 0.02 | 0.035 | 0.003 | 0.086 |
|  |  | feature | 0.018 | 0.021 | 0.397 | -0.024 | 0.06 |
|  | 2 | association | 0.019 | 0.014 | 0.179 | -0.009 | 0.048 |
|  |  | feature | 0.005 | 0.014 | 0.739 | -0.024 | 0.034 |
|  | 3 | association | -0.004 | 0.011 | 0.704 | -0.028 | 0.019 |
|  |  | feature | -0.011 | 0.01 | 0.268 | -0.03 | 0.009 |
|  | 4 | association | 0.01 | 0.01 | 0.296 | -0.009 | 0.03 |
|  |  | feature | 0.009 | 0.014 | 0.523 | -0.02 | 0.038 |
|  | 5 | association | 0.015 | 0.01 | 0.142 | -0.005 | 0.035 |
|  |  | feature | 0.025 | 0.013 | 0.066 | -0.002 | 0.051 |
|  | 6 | association | 0.004 | 0.009 | 0.703 | -0.016 | 0.023 |
|  |  | feature | 0.01 | 0.012 | 0.415 | -0.014 | 0.034 |
|  | 7 | association | .033* | 0.014 | 0.022 | 0.005 | 0.061 |
|  |  | feature | .054* | 0.014 | 0.001 | 0.024 | 0.083 |
|  | 8 | association | .064* | 0.014 | 0 | 0.036 | 0.092 |
|  |  | feature | .088* | 0.012 | 0 | 0.062 | 0.113 |
|  | 9 | association | .113* | 0.019 | 0 | 0.074 | 0.152 |
|  |  | feature | .141* | 0.015 | 0 | 0.11 | 0.172 |
|  | 10 | association | .133* | 0.032 | 0 | 0.066 | 0.199 |
|  |  | feature | .159* | 0.029 | 0 | 0.1 | 0.219 |
| Study 2 context versus emotion | 1 | emotion | 0.016 | 0.02 | 0.432 | -0.025 | 0.058 |
|  |  | context | 0.019 | 0.022 | 0.399 | -0.026 | 0.064 |
|  | 2 | emotion | 0.019 | 0.014 | 0.176 | -0.009 | 0.048 |
|  |  | context | 0.024 | 0.014 | 0.101 | -0.005 | 0.053 |
|  | 3 | emotion | .036* | 0.012 | 0.004 | 0.012 | 0.061 |
|  |  | context | .027* | 0.013 | 0.046 | 0.001 | 0.054 |
|  | 4 | emotion | .073* | 0.014 | 0 | 0.045 | 0.101 |
|  |  | context | .083* | 0.017 | 0 | 0.049 | 0.118 |
|  | 5 | emotion | .078* | 0.018 | 0 | 0.041 | 0.115 |
|  |  | context | .111* | 0.02 | 0 | 0.07 | 0.152 |
|  | 6 | emotion | .062* | 0.015 | 0 | 0.032 | 0.093 |
|  |  | context | .095* | 0.016 | 0 | 0.062 | 0.127 |
|  | 7 | emotion | .050* | 0.016 | 0.003 | 0.018 | 0.082 |
|  |  | context | .070* | 0.013 | 0 | 0.044 | 0.097 |
|  | 8 | emotion | 0.008 | 0.014 | 0.593 | -0.021 | 0.037 |
|  |  | context | 0.024 | 0.014 | 0.113 | -0.006 | 0.053 |
|  | 9 | emotion | -.043* | 0.013 | 0.002 | -0.069 | -0.017 |
|  |  | context | -.035* | 0.014 | 0.017 | -0.064 | -0.007 |
|  | 10 | emotion | -.134* | 0.018 | 0 | -0.171 | -0.097 |
|  |  | context | -.105* | 0.02 | 0 | -0.147 | -0.064 |

Based on estimated marginal means; * The mean difference is significant at the .05 level; ^b^ Adjustment for multiple comparisons: Bonferroni.
